# Motor planning and execution establish distinct feedforward and feedback motor histories

**DOI:** 10.64898/2026.08.27.747554

**Authors:** Christian Seegelke, Tobias Heed

**Affiliations:** Cognitive Psychology, Department of Psychology, University of Salzburg, Austria; Centre for Cognitive Neuroscience, University of Salzburg, Austria

**Keywords:** motor history, reaching, feedback, motor planning, sequential effects, hysteresis, repetition effects

## Abstract

Movements are systematically affected by the recent motor history. These history effects may be induced either by reused motor plans or from lingering tuning of the previous movements’ execution. We dissociated planning and execution using four experimental manipulations across two complementary motor paradigms. We isolated planning by preventing execution with stop signals and mechanical blocks, and execution by moving participants’ hand passively using a robot manipulandum. History effects emerged in feedforward movement aspects – reaction time and early movement kinematics – following isolated planning. In contrast, they were absent or markedly reduced for isolated execution. History effects emerged also in late movement aspect that involves sensory feedback during execution – movement accuracy and precision – but only when movements were both planned and executed. Feedforward effects generalized across hands, whereas feedback effects were effector specific. Thus, prior motor planning and execution make distinct and complementary contributions in shaping future motor behavior.

## Introduction

Recent movements systematically bias subsequent motor behavior^1–9^. Such history dependence implies that information from prior actions is retained and incorporated into the planning and execution of future movements. Characterizing history effects therefore offers leverage to probe the organization and hierarchy of the motor system.

Strikingly, robust history effects can arise after a single movement and manifest in distinct and complementary aspects of motor behavior^10–12^. In unimanual reaching tasks, history primarily biases *how* movements are performed. For example, reaching around an obstacle biases subsequent reaches toward similarly curved trajectories, even after the obstacle is removed, revealing a history-dependent bias in movement kinematics^13,14^. In effector-selection tasks, history primarily affects *how fast* the movement is initiated and biases *which* effector is chosen to perform a movement. For instance, using the same effector twice produces faster responses for the second movement even if movement kinematics differ^6,11,12,15,16^.

The mechanisms underlying history effects remain unresolved. One possibility is that they reflect the recycling of motor plans^7,10,15,17^. According to this account, history effects should arise whenever a movement is planned, even if it is never executed. An alternative possibility is that history effects reflect lingering tuning changes linked to the movement’s execution^3,5,18–20^. According to this account, history effects should emerge whenever a movement was executed, whether it was planned or not.

Discriminating between these two possibilities has been challenging because planning and execution normally co-occur. Here, we systematically dissociate the two aspects in a series of experiments that employ both unimanual and effector-selection paradigms. We used four manipulations that isolate planning by eliminating execution, isolate execution by eliminating planning, or eliminate both (Fig. 1). We used stop signals and mechanical stops to preserve planning while preventing execution, robot-imposed passive movements to induce execution without prior planning, and no-go trials to eliminate both planning and execution. We contrasted these conditions with regularly executed movements that included both planning and execution. In all experiments, we paired two movements – a prime and a probe – such that the prime established the movement history for the subsequent probe. The probe was held constant so that any bias could be attributed to the prime’s planning and execution state.

**Figure 1.**
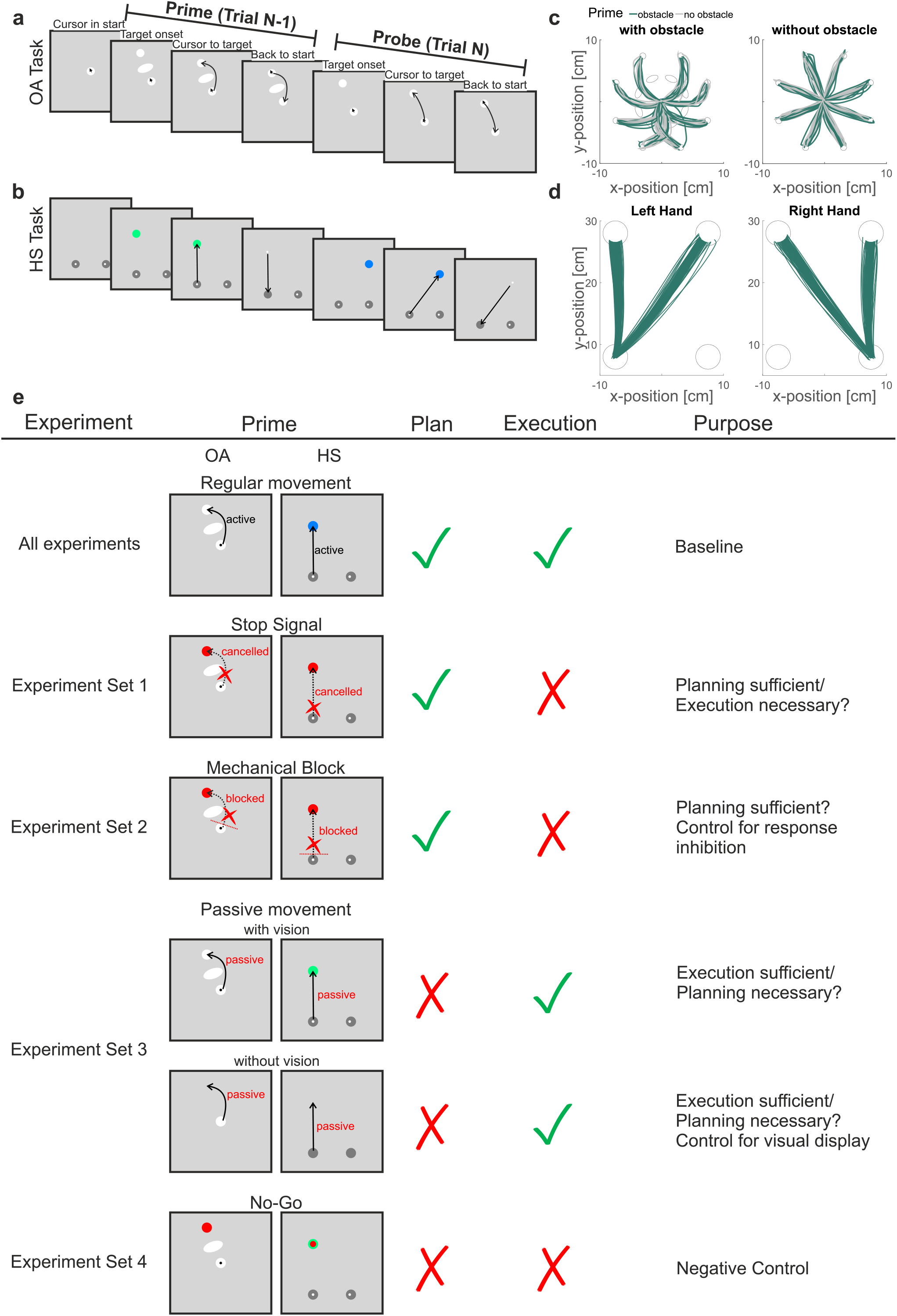
Task procedure and logic of experimental manipulations. Participants performed two consecutive movements (prime and probe, respectively) and performance of probe movement was assessed in dependence of the prime’s characteristics. In the obstacle avoidance (OA) task (**a**), participants performed center-out reaching movements to one of eight targets while avoiding obstacles that occasionally blocked the direct path. In the depicted example, an obstacle was present during the prime phase, necessitating a curved movement while no obstacle was present during the probe phase. In the hand selection (HS) task (**b**), participants performed a reaching movement to one of two targets with either hand; a color cue instructed which hand to use. The example shows a left-hand movement to the left target in the prime phase and a left-hand movement to the right target in the probe phase. **cd** Reach trajectories of probe movements from exemplary participants during active trials in the OA task (**c**) and HS task (**d**). **e** Summary of the experimental logic and applied manipulation to systematically dissociate motor planning and execution of prime movements.

If planning and execution make distinct contributions to history effects, these contributions may be expressed at different stages of motor control. We first examined whether history effects influence feedforward aspects of the movement, which cannot yet be modified by sensory feedback. We assessed reaction time and initial reach direction as feedforward movement aspects. We complemented the feedforward measures with analyses of movement accuracy and precision as later movement characteristics that are susceptible to online corrections based on sensory feedback.

If planning is sufficient for a history effect to emerge, that effect should survive when execution is prevented; if execution is necessary, the effect should instead depend on whether the prime movement was actually carried out or not. We applied this logic separately to feedforward and feedback measures. This analysis approach allows testing whether history effects can emerge at different stages of the motor control hierarchy – planning vs. execution –, and whether the different stages create distinct histories for feedforward- and feedback-related movement aspects.

We report here that movement planning is both necessary and sufficient for the occurrence of history effects on feedforward aspects of a movement. In contrast, combined movement planning and execution were necessary for feedback-related history effects to emerge. The close correspondence of results across the unimanual obstacle-avoidance and the bimanual effector-selection paradigms provides converging evidence that prior motor planning and execution make distinct and complementary contributions in shaping future motor behavior.

## Results

### Feedforward-related history effects are reduced when participants actively inhibit movement

We first confirmed that both tasks exhibited history effects under standard conditions, that is, with primes both planned and executed. For the obstacle avoidance (OA) task (Fig. 1a), the expected history effect was quantified as the difference in initial reach error between probe movements for which the prime movement had, vs. had not, involved an obstacle (i.e. initial reach bias; see Methods). Probe trajectories were more curved after obstacle primes than after unobstructed primes (initial reach bias = 7.9° [6.4, 9.4], brackets contain 95% highest density interval, pd > 99%, 0% in ROPE, see Fig. 2a; we report the estimated marginal means for feedforward measures across the eight main experiments in Supplementary Fig. 1).

**Figure 2.**
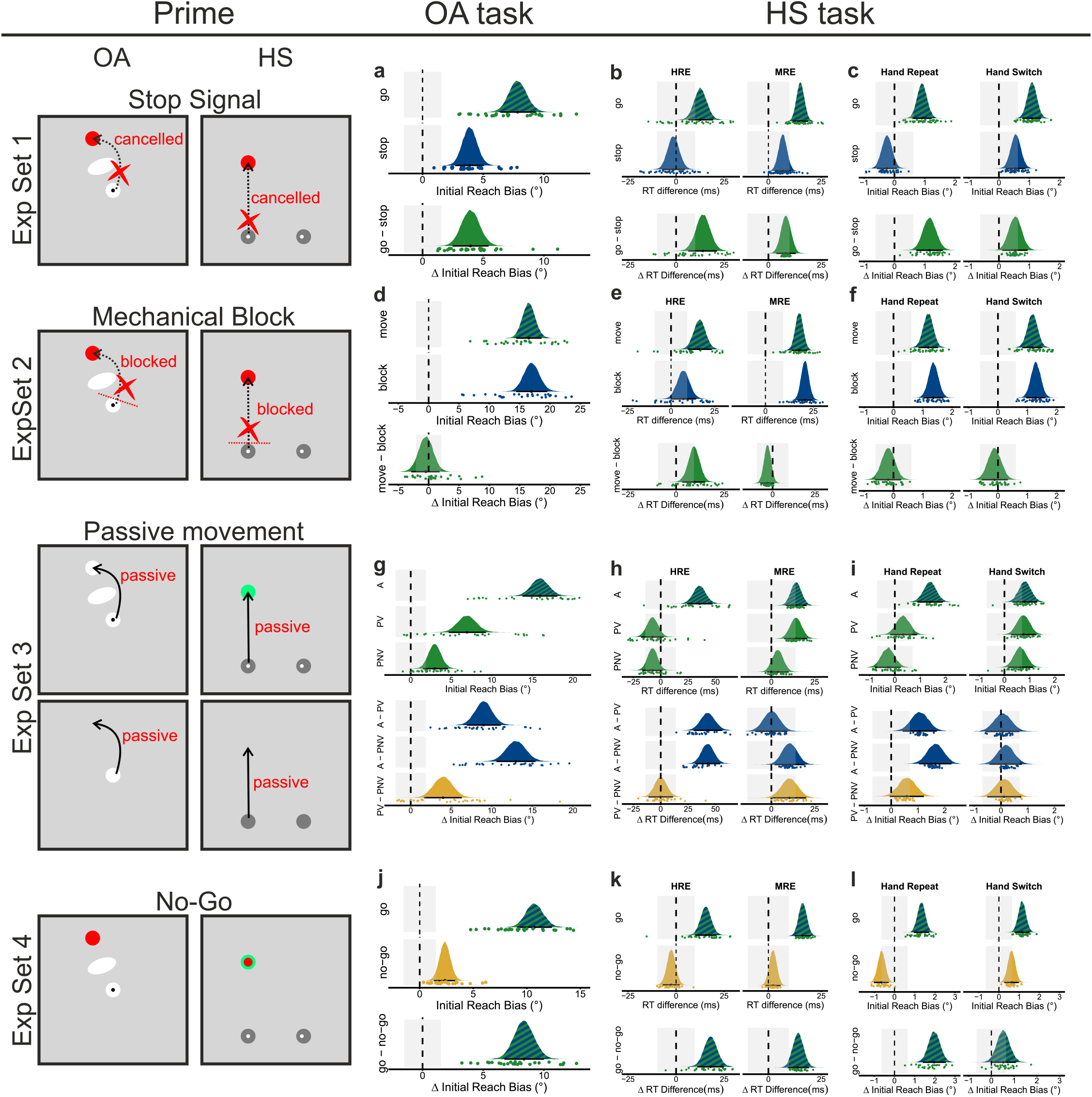
Motor history effects in feedforward measures for the four experimental manipulations (stop signal, mechanical block, passive movements, no-go) across the two tasks. For the obstacle avoidance (OA) task, the history effect was quantified as the difference in initial reach error between probe movements for which the prime movement had, vs. had not, involved an obstacle (i.e. initial reach bias). For the hand selection (HS) task, history effects were quantified as the difference in RT between trials for which 1) different vs the same hand was used for prime and probe movements (hand repetition effect, HRE) and 2) movements were performed in different vs same directions (coded in an egocentric reference frame; movement repetition effect, MRE). Further, we quantified history effects in kinematics (i.e., initial reach bias) as difference in initial reach error between probe movements for diagonal and straight prime movements. Positive values indicate that probe movements were more inclined towards the body-midline following diagonal than straight prime movements. Colored areas: posterior distributions. Black dots: median; error bars: 95% highest-density interval (HDI); grey shaded areas: region of practical equivalence (ROPE). A = active, PV = passive vision, PNV = passive no vision.

For the hand selection (HS) task (Fig. 1b), responses were faster when the probe was executed with the same hand as the prime (hand repetition effect [HRE]: 13 ms [6, 19], pd > 99%, 30% in ROPE) or in the same movement direction (coded in an egocentric reference frame, see Methods; movement repetition effect [MRE]: 15 ms [12, 19], pd > 99%, 0% in ROPE, Fig. 2b). Initial reach direction was also biased towards the preceding prime movement direction (initial reach bias quantified as difference in initial reach error between probe movements for diagonal vs. straight prime movements; positive values indicate that probe movements were more inclined towards the body-midline following diagonal vs. straight prime movements; see Methods). This direction effect generalized to trials in which prime and probe involved different hands (Hand Repeat: initial reach bias = 0.89° [0.58, 1.22], pd > 99%, 7% in ROPE; Hand Switch: 1.09° [0.79, 1.40], pd > 99%, 0% of HDI in ROPE, Fig. 2c). In sum, history effects emerged robustly and were non-negligible in magnitude in our experiments, replicating previous reports^10–12,14^.

Next, we isolated the contribution of movement planning by instructing participants to prepare but cancel execution of the prime movement via a visual stop signal, presented 150ms after prime target onset in a random subset of trials (stop trial). A time constraint on regular go trials discouraged strategic omission of motor planning. For the OA task, a reliable history effect was evident in stop trials (initial reach bias = 3.9° [2.7, 5.1], pd > 99%, 0% in ROPE, Fig. 2a), albeit reduced in size (difference go vs stop = 4.0° [2.5, 5.5], pd > 99%, 0% in ROPE, Fig. 2a). Thus, planning alone elicited a motor history effect on a feedforward-related kinematic measure.

In the HS task, the stop signal strongly reduced history effects. The HRE was absent (-1 ms [-7, 5], pd = 68%, 100% in ROPE; difference go vs stop: HRE: 14 ms [6, 21], pd > 99%, 24% in ROPE) whereas the MRE was present but reduced in magnitude (7 ms [3, 11], pd > 99%, 91% in ROPE, Fig. 2b; difference go vs stop: 8 ms [4, 13], pd > 99%, 67% in ROPE, Fig. 2b). Similarly, initial reach bias was absent or attenuated (Hand Repeat: -0.25° [-0.59, 0.08], pd = 93%, 100% in ROPE; Hand Switch: 0.56° [0.23,0.88], pd > 99%, 60% in ROPE, Fig. 2c; difference go vs stop Hand Repeat: 1.16° [0.72, 1.60], pd > 99%, 0% in ROPE, Hand Switch: 0.54° [0.11, 0.95], pd > 99%, 61% in ROPE).

Together, Experiment Set 1 indicates that isolated planning is sufficient to establish history effects on feedforward-related measures, but these effects were markedly reduced, and partly eliminated, relative to plan-and-execute trials.

### Feedforward-related history effects persist when execution is physically prevented

Because stop-signal trials recruit response inhibition^21,22^, the size reduction of history effects could reflect either a contribution of movement execution to history effects or the suppression of preparatory activity when an ongoing task must be stopped^23,24^. To distinguish these possibilities, Experiment Set 2 physically blocked the prime movement while preserving motor planning (blocked trials); physical blocking removes the need for participants to inhibit the movement themselves. In the OA task, mechanically blocking prime execution did not alter history effects. Initial reach bias was comparable between unblocked trials (i.e., move trials: 16.5° [14.4, 18.6], pd > 99%, 0% in ROPE) and blocked trials (17.0° [14.4, 19.6], pd > 99%, 0% in ROPE; difference move vs block: -0.5° [-2.9,1.8], pd = 65%, 83% in ROPE, Fig. 2d).

In the HS task, too, mechanical blocking had little impact on history effects. The HRE was present but reduced (7 ms, [0,14], pd > 97%, 64% in ROPE; difference move vs block: 9 ms, CI [3,15], pd > 99%, 51% in ROPE), whereas the MRE remained fully intact (21 ms, CI [17,25], pd > 99%, 0% in ROPE; difference move vs block: -3 ms [-7, 1], pd = 92%, 100% in ROPE, Fig.2e). Initial reach bias was present and unchanged for both repeated and switched hands (blocked trials Hand Repeat: 1.36° [1.03,1.70], pd > 99%, 0% in ROPE, Hand Switch: 1.29° [0.96, 1.62], pd > 99%, 0% in ROPE; difference move vs block Hand Repeat: -0.18° [-0.62, 0.26], pd = 79%, 99% in ROPE, Hand Switch: -0.12° [-0.59, 0.33], pd = 70%, 100% in ROPE], Fig. 2f).

In sum, feedforward-related history effects were largely unaffected when execution was physically prevented. Accordingly, the reduction of history effects observed in Experiment Set 1 was likely due to the cognitive requirement of response inhibition rather than the absence of execution alone. A decisive test of necessity is therefore whether history effects emerge when movements are executed without prior planning — that is, whether planning is necessary for establishing motor history.

### Motor execution without prior planning largely obliterates feedforward-related history effects

In Experiment Set 3, the prime consisted of robot-imposed, passive movements that replayed each participant’s previously recorded, active trajectories. Accordingly, active and passive prime movements exhibited highly similar spatiotemporal characteristics, and we ascertained that participants did not, against the instructions to be guided passively, exert force in movement direction (Supplementary Fig. 2).

In the OA task, passive primes evoked a reduced but reliable history effect (initial reach bias: 6.9° [4.7, 9.3], pd > 99%, 0% in ROPE, Fig. 2g; difference of active vs. passive vision: 9.0° [6.9, 11.1], pd > 99%, 0% in ROPE). To test whether the visual display may induce the remaining history effect, we included a condition in which we withheld all visual information during passive primes (no vision); the bias was further reduced but remained reliable (passive no vision: 3.0° [1.5, 4.6], pd > 99%, 12% in ROPE; difference active vs. passive no vision: 13.0° [10.6, 15.4], pd > 99%, 0% in ROPE; difference passive vision vs. passive no vision: 4.0° [1.6, 6.3], pd > 99%, 5% in ROPE, Fig. 2g). Finally, we ran an active task in which we withheld the visual display during prime movements (see active movements with and without visual feedback in Methods for details).

This did not affect history effects compared to the regular active task with visual prime display (difference vision vs no-vision during active primes: -0.1° [-2.1, 1.9], pd = 53%, 99% in ROPE, Supplementary Fig. 3). Thus, the reduction of history effects after passive primes was due to the absence of planning, not the absence of a visual display.

In the HS task, passive primes largely eliminated all history effects. The HRE was absent regardless of visual display (with vision: -8 ms [-18, 3], pd = 93%, 81% in ROPE; no vision: -7 ms [-17, 2], pd = 95%, 85% in ROPE; difference active vs passive vision: 43ms [30, 56], pd > 99%, 0% in ROPE; active vs passive no vision: 43ms [32, 54], pd > 99%, 0% in ROPE, Fig. 2h). A residual MRE persisted only when visual information was available, suggesting that this component may partly reflect non-motor contextual or visuospatial repetition effects (with vision: 14 ms [7, 21], pd > 99%, 50% in ROPE; no-vision: 4 ms [−3, 10], pd = 88%, 100% in ROPE; difference active vs passive vision: 0ms [-9, 9], pd = 51%, 100% in ROPE; active vs passive no vision: 11ms [2, 19], pd > 99%, 70% in ROPE, Fig. 2h). Probe kinematics mirrored this pattern: active primes yielded an initial reach bias for both Hand Repeat (1.39° [0.89, 1.89], pd > 99%, 0% in ROPE) and Hand Switch conditions (0.81° [0.32, 1.30], pd > 99%, 39% in ROPE, Fig. 2i); passive primes yielded no bias when the hand was repeated (with vison: 0.32° [-0.18, 0.87], pd = 88%, 84% in ROPE; without vision: -0.27° [-0.79,0.23], pd = 85%, 91% in ROPE; difference active vs vision: 1.06° [0.43, 1.70], pd > 99%, 21% in ROPE; active vs no vision: 1.66° [1.05, 2.25], pd > 99%, 0% in ROPE, Fig.2i) but a reliable (albeit small) bias when the hand switched, irrespective of vision (with vision: 0.75° [0.26, 1.27], pd > 99%, 44% in ROPE; without vision: 0.63° [0.15,1.13], pd > 99%, 56% in ROPE; difference active vs vision: 0.07° [-0.58, 0.69], pd = 58%, 100% in ROPE; active vs no vision: 0.18° [-0.43, 0.78], pd = 72%, 93% in ROPE, Fig. 2i).

Together, these results show that planning is necessary for robust feedforward history effects: execution without planning failed to produce the HRE and substantially reduced kinematic biases. However, the residual effects observed after passive primes, at least when the visual display is available, suggest that execution may make a small additional contribution. Alternatively, these residual biases may reflect incidental planning, motor imagery, or task-level contextual expectations acting as weak priors on subsequent movements^14^.

### Residual history effects remain even if both motor planning and execution are eliminated

If the residual biases observed after passive primes arise from movement execution, then removing both planning and execution should further reduce history effects. In Experiment Set 4 we therefore used a go/no-go manipulation with a 50/50 ratio for primes: the visual display occurred, and an immediate cue indicated whether participants should move or withhold movement.

In the OA task, we first interleaved no-go and go trials within blocks. A small but reliable residual reach bias remained on no-go trials (2.4° [1.4, 3.3], pd > 99%, 10% in ROPE), though it was substantially reduced relative to go trials (difference go vs no-go: 8.4° [6.8, 10.0]; pd > 99%, 0% in ROPE, Fig. 2j). Therefore, we tested whether random trial interleaving promoted covert preparation because participants expected that they might have to move. We presented go and no-go trials in separate blocks, so that participants could effectively disregard the prime display in no-go blocks. Still, a qualitatively similar bias persisted, arguing against uncertainty-induced covert planning as the primary source of the residual bias (Supplementary Fig. 4). Because no-go trials produced a small residual bias in the absence of both planning and execution, we reasoned that the difference between go and no-go trials, which eliminates the unrelated residual bias, can serve as an estimate of the combined contribution of motor planning and (potentially) execution to history effects. We then subtracted difference scores for each experimental manipulation – go minus stop (Experiment 1), move minus block (Experiment 2), and active minus passive (with and without visual feedback; Experiment 3) – from the go–no-go difference score. Each experimental difference score isolates either planning or execution, so that the subtraction estimates the contribution of the respective other process.

Accordingly, subtracting the execution-related difference scores in Experiments 1 and 2 from the go– no-go difference yielded an estimate of the contribution of planning, whereas subtracting the planning-related difference scores in Experiment 3 yielded an estimate of the contribution of execution. Positive values indicate a residual bias attributable to the isolated process.

The reduction in reach bias was larger under the no-go condition (Experiment Set 4) than under both the delayed-stop condition (Experiment Set 1; difference: 4.4° [2.2, 6.6], pd > 99%, 0% in ROPE) and the mechanical-block condition (Experiment Set 2; difference: 8.8° [6.0, 11.7], pd > 99%, 0% in ROPE).

This pattern suggests that history effects are attributable primarily to motor planning. In contrast, the reduction in reach bias was comparable to that observed after passive movement with visual feedback (Experiment Set 3; difference: −0.6° [−3.3, 2.0], pd = 68%, 65.8% in ROPE) and was in the opposite direction relative to passive movement without visual feedback (difference: −4.6° [−7.5, −1.7], pd > 99%, 1% in ROPE). Thus, prior movement execution appears to contribute little, if anything, to the residual bias.

In the HS task, both HRE and MRE were absent after no-go primes (HRE: -3 ms [-7, 2], pd = 87%, 100% in ROPE; difference go vs no-go: 18ms [11,25], pd > 99%, 0% in ROPE; MRE: 2 ms [-1, 6], pd = 89%, 100% in ROPE; difference go vs no-go: 14ms [9,19], pd > 99%, 6% in ROPE; Fig. 2k). Furthermore, go primes yielded an initial reach bias for both Hand Repeat (1.33° [0.96, 1.72], pd > 99%, 0% in ROPE) and Hand Switch conditions (1.15° [0.77, 1.51], pd > 99%, 0% in ROPE); no-go primes yielded a small opposite bias when the hand was repeated (-0.64° [-1.01, -0.29], pd > 99%, 50% in ROPE; difference go vs no-go: 1.97° [1.40, 2.55], pd > 99%, 0% in ROPE) and a small bias when the hand switched (0.63° [0.27, 0.98], pd > 99%, 54% in ROPE; difference go vs no-go: 0.52° [-0.03, 1.08], pd = 97%, 61% in ROPE, Fig. 2l).

In sum, results from using passive primes (execution, no planning) and no-go primes (no execution, no planning) support the same conclusion: robust feedforward history effects require planning. The small residual biases that survive the experimental elimination of planning and execution—especially in OA—are best attributed to task-level context (e.g., sustained expectations about obstacle-induced curvature), not to execution-dependent carryover.

### Planning and execution are necessary for the emergence of history effects on feedback-related movement aspects

So far, we have shown that motor planning is both necessary and sufficient to bias feedforward aspects of subsequent movements. We next examined whether history effects extend to feedback-related aspects of movement control, quantified as the difference in absolute reach endpoint errors (i.e., reach endpoint accuracy) and reach endpoint variability (i.e., reach endpoint precision) between trials with different versus same prime–probe movement requirements (OA: obstacle presence/ absence; HS: different vs. same movement direction; positive values indicate better performance for same prime–probe movement requirements; see Methods).

In the OA task, prime movements that were planned and executed produced a reliable (but negligible in magnitude) difference in probe reach accuracy (pooled across the four experiments: 0.04mm [0.02, 0.07], pd > 99%, 100% in ROPE, Fig. 3aceg, we report the estimated marginal means for all feedback measures across the eight main experiments in Supplementary Fig. 7). This pattern is consistent with prior work showing that active movement generation creates a small motor history for feedback-related movement aspects^14^. Critically, this modest evidence of an effect was present only in the fully planned-and-executed condition; when execution was cancelled via a delayed stop-signal, existence itself was no longer supported (Exp. 1: 0.01mm [-0.07, 0.05], pd = 66%, 100% in ROPE, Fig. 3a), nor when execution was mechanically blocked while the plan remained intact (Exp. 2: - 0.01mm [-0.06, 0.05], pd = 61%, 100% in ROPE, Fig. 3c). Robot-imposed passive primes (Exp. 3) likewise gave no evidence that an effect existed, regardless of whether the visual display was shown (vision: 0.04mm [-0.02, 0.08], pd = 90%, 100% in ROPE; no vision: 0.03mm [-0.03, 0.08], pd = 86%, 100% in ROPE, Fig. 3e), as did primes with neither planning nor execution (Exp. 4: 0.01mm [-0.04, 0.06], pd = 64%, 100% in ROPE, Fig. 3g). A similar pattern emerged for end-point precision and final reach error (Supplementary Fig. 5-7). Viewed together, the evidence for a feedback-related history effect in OA was strongest — though still negligible in absolute size — only when planning and execution co-occurred, whereas none of the isolated-process conditions provided comparable evidence that an effect existed at all. This pattern contrasts with that of feedforward aspects, where planning alone was sufficient to produce effects that both credibly existed and were, in most cases, non-negligible in magnitude.

**Figure 3.**
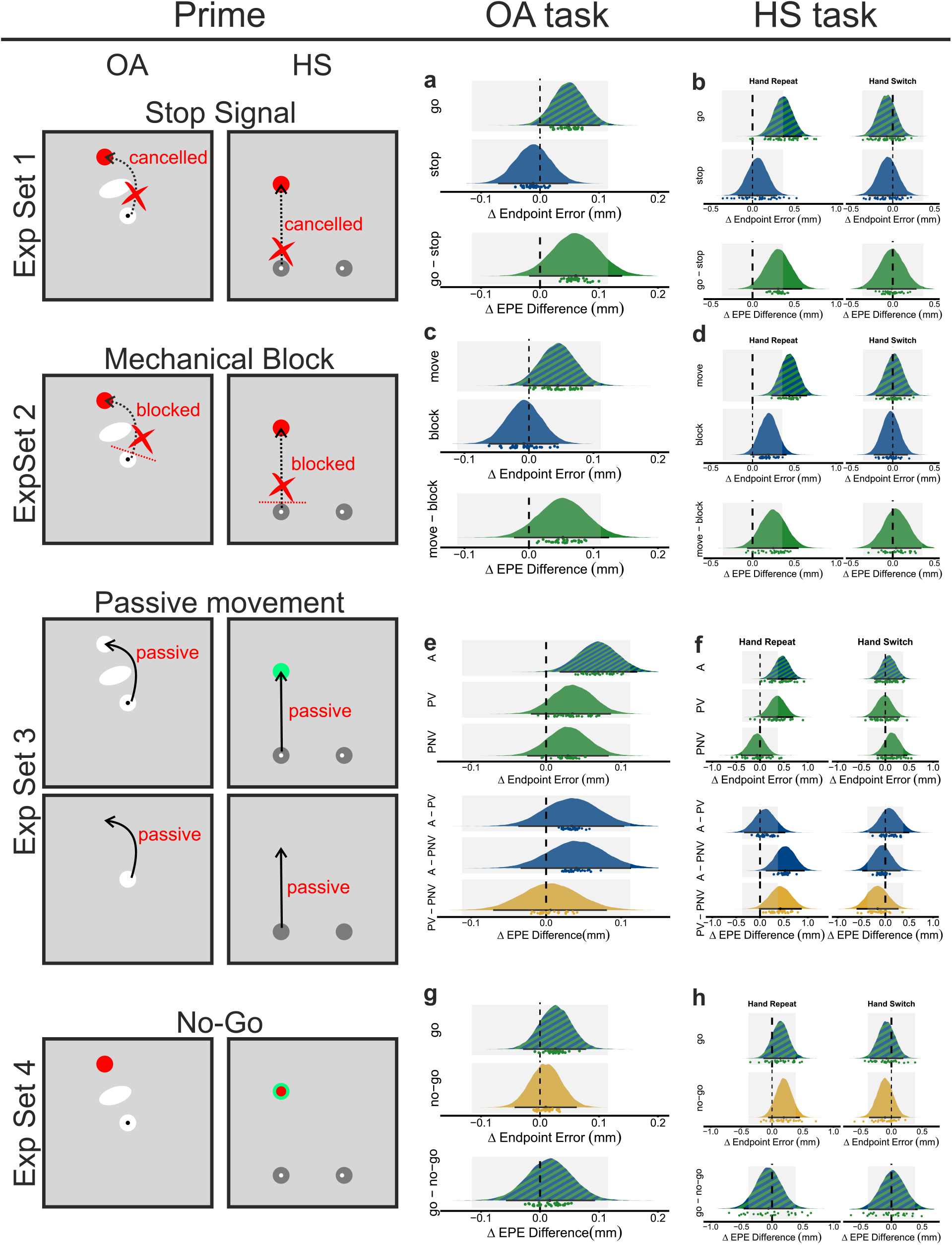
Motor history effects in feedback measures (reach endpoint accuracy) for the four experimental manipulations (stop signal, mechanical block, passive movements, no-go) across the two tasks. History effects were quantified as difference in absolute reach endpoint error (in mm) between trials with different versus same prime–probe movement requirements (OA: obstacle presence/ absence; HS: different vs. same movement direction). Positive values indicate better performance (i.e., smaller endpoint errors) for same prime–probe movement requirements. Colored areas: posterior distributions. Black dots: median; error bars: 95% highest-density interval (HDI); grey shaded areas: region of practical equivalence (ROPE). A = active, PV = passive vision, PNV = passive no vision.

In the HS task, the improvement in reach accuracy for matched prime–probe movement requirements during active trials likely existed when the same hand was used across prime and probe (pooled across the four experiments: 0.35mm [0.25, 0.47], pd > 99%), and, unlike the OA effect, its magnitude was at least partly outside the ROPE (43% of the HDI outside the region), providing some evidence that this effect was non-negligible. No comparable evidence of an effect emerged when hands switched (-0.03mm [-0.14, 0.09], pd = 69%, 100% in ROPE, Fig. 3bdfh). As in OA, cancelling a planned movement after a stop-signal (Exp. 1) or mechanical blocking (Exp. 2) removed evidence for the effect’s existence entirely (Exp. 1: hand repeat: 0.06mm [-0.18, 0.30], pd = 69%, 100% in ROPE; hand switch: -0.06mm [-0.30, 0.17], pd = 69%, 100% in ROPE, Fig. 3b; Exp. 2: hand repeat: 0.19mm [-0.03, 0.41], pd = 96%, 88% in ROPE; hand switch: -0.03mm [-0.23, 0.18], pd = 60%, 100% in ROPE, Fig. 3d), diverging from the feedforward pattern, where isolated planning produced a reliable effect. Passive primes (Exp. 3) showed a similar profile. With vision present, the hand-repeat effect likely existed but its magnitude was inconclusive, (hand repeat: 0.36mm [0.02, 0.70], pd = 98%, 53% in ROPE, Fig. 3f); the hand-switch effect showed no credible evidence of existing (-0.02mm [-0.35, 0.31], pd = 54%, 100% in ROPE). Without vision, neither hand condition showed credible evidence of an effect (hand repeat: -0.06mm [-0.40, 0.27], pd = 65%, 97% in ROPE; hand switch: 0.14mm [-0.18, 0.46], pd = 81%, 87% in ROPE). Likewise, there was no evidence for an effect when both planning and execution were absent (Exp. 4: hand repeat: 0.19mm [-0.06, 0.45], pd = 93%, 87% in ROPE; hand switch: -0.10mm [-0.36, 0.15], pd = 79%, 100% in ROPE, Fig. 3h). Endpoint precision and final reach error again followed the same pattern (Supplementary Fig. 5-7). Thus, as in OA, the result pattern suggests planning and execution jointly contribute to feedback-related history effects in HS, and the effect that does emerge is effector-specific.

Across tasks, evidence for feedback-related history effects was present — though still small — when movements were both planned and executed; when either process was isolated, there was no credible evidence that an effect existed at all. This pattern thus diverges from the feedforward findings and suggests that the joint occurrence of planning and execution provides the most consistent basis for feedback-related history effects, whereas planning or execution alone does not reliably establish them.

## Discussion

Movements that occur earlier in a sequence leave a systematic imprint on subsequent motor performance^3–5,9–14,16^. Here, we systematically dissociated the contributions of prior planning and execution in shaping these history effects.

Three principal findings emerged. First, motor planning, but not execution, was necessary and sufficient for history effects on feedforward movement aspects to occur, expressed reliably in early movement kinematics and response latency. Second, history effects also existed, albeit small in magnitude, for feedback-related movement aspects; however, they only emerged when movements had been planned and executed.

Third, history effects on feedforward movement aspects transferred across hands, indicating that some elements of the motor history are effector-independent. Again, the result pattern diverged for feedback-related movement aspects, for which no generalization across effectors was evident. Thus, history effects appear tied to the effector here. The disparities between feedforward and feedback-related movement aspects point to distinct contributions of motor planning and execution in shaping subsequent movements.

Our findings map well onto a dynamical systems account for motor cortex, which emphasizes the role of motor planning^25,26^ for the evolution of subsequent motor behavior. In this framework, motor planning organizes the neural population into an initial state in population activity space, from which movement-generating dynamics subsequently unfold^25–28^. Different motor plans therefore place the neural population into distinct initial states within the preparatory subspace, giving rise to individual neural trajectories that evolve dynamically into specific motor output patterns. In this view, feedforward movement aspects (i.e., early movement kinematics) depend largely on the network’s state at the time movement execution is triggered, with subsequent motor-related activity emerging from the ensuing neural dynamics rather than independently specifying movement parameters.

This framework has also received support from behavioral experiments in humans. If distinct motor plans map to distinct preparatory neural states, then identical movements can be associated with different motor memories provided they are preceded by different plans. Consistent with this prediction, planning (but never executing) two different follow-through movements allows humans to adapt to two different force fields, which would otherwise interfere and obstruct adaptation^29^. Similarly, motor learning can occur through movement preparation even without overt movement execution^30^.

History effects in feedforward control integrate naturally into this framework. If motor planning establishes an initial preparatory state, then trial-to-trial history effects can arise when the network does not fully reset between successive movements. Instead, residual activity from the previous trial can bias the preparatory state from which subsequent movement-generating dynamics unfold, thereby influencing early movement kinematics. Similarly, accelerated reaction time may reflect history-dependent activity in the neural state that monitors the transition from preparation to movement execution, consistent with evidence that movement timing is represented by population dynamics distinct from those specifying movement identity^31^. Thus, residual activity from one trial may influence both the trajectory of the ensuing movement and the timing with which those dynamics are engaged on the subsequent trial.

This framework also provides a principled explanation for why the stop-signal condition produced a reduced, rather than abolished, history effect, whereas the mechanical-block condition preserved it. Successfully cancelling a planned movement, unlike being physically prevented from executing it, requires active engagement of the fronto-basal ganglia action-stopping network^32–34^. This inhibitory process is thought to suppress or disrupt the evolving preparatory state, rather than merely preventing its downstream motor output^23,35,36^. One possibility is that successful stopping partially reshapes the preparatory state, reducing—but not eliminating— the extent to which it can influence subsequent trials.

Conversely, when movement execution is physically blocked after the motor command has been generated, the preparatory state may remain largely intact, given that the movement has been triggered and only failed to execute properly. Thus, its residual planning-related neural activity may persist into the next trial. While these interpretations remain to be tested directly with neural population recordings, they provide a mechanistic account of why active stopping and passive movement blockage produce distinct effects on trial-to-trial history.

The dissociation between feedforward and feedback history effects further suggests that prior actions leave distinct traces within the motor system. Only movements that had been planned and executed produced a feedback-stage bias, whereas planning alone did not. Moreover, passive execution in the absence of prior planning was insufficient to produce comparable feedback effects. This pattern suggests that feedback-related history effects do not simply reflect sensory consequences of prior movement but instead depend on the persistence of a planned motor state and the predictive processes engaged during preparation.

This distinction bears resemblance to classic repetition-effect models in the priming literature, which have distinguished between facilitation and sharpening mechanisms^37–39^. Facilitation models propose that repetition reduces processing demands by increasing the efficiency or accessibility of a previously established representation, without necessarily altering its underlying structure. This framework provides a useful analogy for our feedforward history effects: maintaining the previous preparatory state may allow the motor system to readily re-enter previously occupied regions of preparatory state space, reducing the cost of establishing the initial conditions from which movement-generating dynamics unfold. In this sense, feedforward history effects may reflect a facilitation-like process whereby prior planning biases subsequent motor trajectories through persistent or partially reactivated preparatory states.

By contrast, sharpening models propose that repetition enhances the precision or selectivity of the neural representation supporting a behavior. This account provides a closer analogy for feedback-related history effects. Final movement accuracy depends on online state estimation and correction processes, in which predicted limb states are compared with sensory feedback to update motor commands^40–43^. Increasing the precision of a predictive estimate for the upcoming movement would require generating and retaining a specific motor prediction (e.g., via efference copy) and comparing that prediction to incoming sensory feedback to update control. Consistent with this interpretation, feedback-related history effects did not emerge when execution was prevented (stop signal or mechanical block) or when execution occurred without prior planning (passive movement) but were observed only when the movement was both planned and executed; thus, they appear to depend on the perpetuation of an internally generated and implemented motor plan. Feedforward and feedback effects may therefore reflect two different forms of persistence within the motor system: carryover of the initial conditions governing a dynamical trajectory (facilitation-like) versus persistence of the predictive state estimates used for online control (sharpening-like) — the latter requiring, specifically, that the motor plan be generated and maintained without interruption. Another important dissociation concerns the transfer of history effects across effectors: history effects on feedforward movement aspects generalized across the hands, whereas those on feedback-related aspects did not. This pattern aligns with emerging evidence that motor cortex does not represent different effectors through entirely separate circuits but rather organizes information through partially overlapping population-level subspaces. Neural population studies have shown that task- and effector-related variables can be distributed across individual neurons while becoming segregated at the level of population dynamics. For example, individual neurons can be active during motor activity related to both arms, while the population activity associated with each arm nevertheless occupies largely distinct, near-orthogonal subspaces^44^. Within this framework, the transfer of feedforward biases in early movement direction across the hands suggests that the planning-related state that influences subsequent movement generation is supported by a shared, effector-general state. In contrast, the absence of transfer for feedback-related biases suggests that the retained information supporting online correction is embedded within effector-specific dimensions, consistent with the requirement that feedback control estimate the current state of the limb and compare it with the predicted consequences of the motor command^40,41,43^ — processes necessarily coupled to the biomechanics of the specific effector involved. Rather than reflecting a simple abstract-vs-specific dichotomy, then, motor history effects appear organized by the computational role of the retained state — generalizable for planning, effector-partitioned for control — thus providing a behavioral signature of the same organizational principle observed in motor cortical population dynamics: the coexistence of shared computational dimensions and effector-specific subspaces within a common neural population.

Taken together, our findings support a unified framework in which motor history effects arise from the persistence of distinct, planning-related motor states. Feedforward carry-over may reflect the partial reuse of an effector-general preparatory state established during motor planning, biasing subsequent movement initiation and early movement trajectories even when the planned movement is never executed. By contrast, feedback-related carry-over appears to depend on the formation and maintenance of an effector-specific predictive state that supports online state estimation and correction — a state that is established only once a planned movement is actually carried out. Because such predictive representations require not only a preserved motor plan but its actual implementation, feedback-related history effects emerged only when movements were both planned and executed, and were absent whenever execution was prevented — whether the plan was physically blocked, actively cancelled, or bypassed altogether via passive movement without prior planning.

Thus, feedforward and feedback history effects reflect distinct forms of persistence within the motor system: the former arises from reuse of preparatory states, whereas the latter depends on maintenance of predictive motor command.

Within this framework, response inhibition is not simply a mechanism that prevents the downstream expression of a motor command. Instead, successful stopping actively alters the preparatory state from which movement dynamics would otherwise unfold, disrupting the persistence of both planning-related and prediction-related information. This account provides a mechanistic explanation for why stop-signal trials differed from physically blocked trials even though both paradigms abolish overt movement execution: the former erodes the internal motor state, whereas the latter preserves it.

Thus, across the four experimental manipulations, the pattern of results is consistent with a hierarchical organization of motor history effects: Planning establishes a transferable preparatory state that shapes future movement generation and generates a predictive estimate of the movement’s outcome; execution allows this prediction to be compared against actual sensory feedback, forming the effector-specific states that support feedback control; and inhibitory processes determine whether these internal states are preserved or disrupted. Therefore, motor history is not a single, undifferentiated memory of prior actions but arises from the persistence of multiple, interacting processes at different stages of the motor control hierarchy.

## Methods

### Participants

We collected data from a total of 320 individuals across 10 experiments. All participants reported to have normal or correct-to-normal vision and received course credit in exchange for their participation. The University of Salzburg Ethics committee and Bielefeld Ethics Committee approved all procedures (Ethical Application Ref: EK-GZ 32/2023 and 2017-114). We excluded data from participants who exhibited high (>30% in the Obstacle Avoidance [OA]Task or >50% in the Hand Selection [HS] Task) error rates, did not follow task instructions, or did not complete the experiment (Experiment Set 1 OA: N=0; HS: N = 1; Experiment Set 2 OA: N=0; HS: N = 1; Experiment Set 3 OA: N=5; OA Control: N = 0; HS: N = 0; Experiment Set 4 OA: N=1; OA Control: N = 0; HS: N = 1).

### General task setups and procedures

We used two different tasks: 1) an obstacle avoidance (OA) task and 2) a hand selection (HS) task. In Experiment Set 1 and 4, we employed the obstacle avoidance task in a mouse tracking paradigm.

Participants controlled an optical computer mouse (MS116, Dell Inc.) with their dominant hand. Stimuli were presented against a grey (RGB [128 128 128]) background on a 25” computer monitor (AW2523HF, Dell Inc.; 1920×1080 pixels; 240Hz refresh rate) positioned at a viewing distance of about 50 cm.

Stimulus presentation and response recording were controlled with PsychoPy v2023.2.3^45^. In Experiment Set 2 and 3 (OA task) and in all HS tasks, participants sat in front of a robotic manipulandum (Kinarm End-Point Lab) while grasping the handle(s) with their hand(s). Movements were restricted to the horizontal plane and stimuli were projected from a horizontal screen (60Hz refresh rate for Experiment Set 1, 2, and 3 (HS task) and 120Hz refresh rate for Experiment Set 4 (HS task) and Experiment Set 2 and 3 (OA task)) onto an opaque mirror that was positioned halfway between the screen and the handles. The mirror prevented participants from seeing their arms, hands, and handles. Tasks were programmed in MATLAB version 2015a (HS task Experiment Set 1, 2, and 3) or MATLAB version 2019b (HS task Experiment Set 4; OA task Experiment Set 2 and 3) and controlled with Kinarm’s Dexterit-E software version 3.6 (HS task Experiment Set 3), 3.9 (HS task Experiment Set 1 and 2) and 3.10 (HS task Experiment Set 4; OA task Experiment Set 2 and 3). End-point forces at the handles were measured with ATI mini 40 six degree of freedom force-torque transducers (ATI Industrial Automation, Apex, NC, USA). The Kinarm recorded hand position and force data at 1000 Hz.

### Obstacle Avoidance Task

Participants performed two successive movements (a prime and a probe movement, respectively) in which they moved a cursor from a central start location to one of 8 target locations. The target locations were arranged on an imagery circle of 8 cm (Experiment Set 1 and 4) or 10 cm (Experiment Set 2 and 3) from the start location.

They were equidistantly spaced at 45° intervals with their angular positions rotated 22.5° from the cardinal directions. In a subset of trials, an obstacle appeared between start and target location which had to be avoided. Start and target locations were white circles (1 cm in diameter). Obstacles were white ellipses (major axis = 2 cm, minor axis = 1 cm) and positioned halfway between start and target location with the major axis oriented perpendicular to a line connecting start and target location. Of particular interest was the performance of the second (i.e., the probe) movement depending on task characteristics during the preceding (i.e., the prime) movement (i.e., presence/ absence of obstacle and other manipulations, see specific task setups and procedures). Importantly, while the specifics of the prime phase differed across experimental conditions, the probe phase remained invariant such that participants always actively performed the probe movement. This allowed us to attribute any performance difference measured in the probe movement to our manipulation of the preceding prime movement.

### Hand Selection Task

Participants performed two successive reaching movements (a prime and a probe movement, respectively) with their right or left hand to move a white circular cursor (1.0 cm in diameter) from a start to one of two target locations. Two circles (diameter = 4.0 cm) shown at the bottom center of the workspace and spaced 15 cm apart served as start locations for the left and right cursors, respectively. Two circular target locations (diameter = 4.0 cm) were positioned at a vertical distance of 20 cm from the two starting locations. The target locations could be displayed in different colors, and the color indicated which hand to use. Again, of particular interest was the performance of the second (i.e., the probe) movement depending on task characteristics during the preceding (i.e., the prime) movement (i.e., repetition of hand and/or movement direction and other manipulations, see specific task setups and procedures). We again systematically manipulated the specifics of the prime phase across experimental conditions and kept the probe phase invariant such that participants always actively performed the probe movement such that we could attribute performance differences during the probe movement to prime phase manipulation.

### Specific task setups and procedures

#### Experiment Set 1: Stop Signal

##### Obstacle Avoidance

30 right-handed participants (24 female, 6 male; mean age = 22.5 years, SD = 4.5 years; mean EHI score = 84.3, SD = 20.3) controlled a mouse cursor to perform two center-out reaching movements in succession. Each trial (comprised of a prime and a probe movement) started with the presentation of the start location at the center of the screen. Participants then moved the cursor into this location and maintained it there for 500 ms. During go trials, the prime target then appeared at one of the 8 target locations, and participants moved the cursor to the respective target location. After the cursor was maintained at the target location for 250 ms, the target location disappeared, and the participant moved the cursor back to the start location, marking the end of the prime phase. After maintaining the cursor there for 500 ms, the probe target appeared (at the same location as in the prime), and participants moved the cursor to the target location. After the cursor was maintained at the location for 250 ms, the participant then moved the cursor back to the start location triggering initiation of the next trial. In a subset of trials, an obstacle appeared either during the prime phase, the probe phase, or both. If present, the obstacle appeared simultaneously with the target location and participants were instructed to move the cursor around the obstacle to reach the target location. The obstacle remained present also during the back-to-start movements and was extinguished once participants moved the cursor back in the start location. During stop trials, the prime target location turned red (stop-signal) 150 ms after prime target onset for 500 ms, signaling participants to withhold their movement. After participants maintained the cursor at the start location for 2000 ms (to approximately match the duration between prime target onset and probe target onset of go trials), the probe phase was initiated, which was identical for both trial types. During go trials, participants had to move the cursor out of the start position within 750 ms after target onset, otherwise the message “React faster!” was shown on the screen in black letters for 1000 ms. Participants still had to reach the target location and back to the start location to initiate the next phase/ trial. If participant hit the obstacle, it turned red for 500 ms and the message “You shall not pass!” was displayed on the screen in black for 1000 ms. If participants moved the cursor out of the start location during a stop-trials, the message “Stop-Signal: Don’t move!” was displayed on the screen in black for 1000 ms, and there was a delay of 1500 ms before participants could initiate the probe phase.

Participants performed 11 blocks, each consisting of 64 trials. In each trial, we manipulated if an obstacle was present or absent during the prime and/ or probe phase. Half of the trials were go-trials, the other half stop-trials, and movements were to be made to each of 8 the target locations with target location held constant between prime and probe phase. Thus, there were a total of 64 conditions, comprised of the factors 2 obstacle prime (present, absent) x 2 obstacle probe (present, absent) x 2 trial type (go-trial, stop-trial) x 8 target location. In each block, each condition was presented once, and the order of presentation was randomized. The first block was considered practice and not analyzed. In total, the experiment comprised 640 trials (without practice) and took about 1.2 hours.

##### Hand Selection

39 participants (25 female, 14 male; mean age = 25.6 years, SD = 7.6; 33 right-handers: mean EHI score = 81.3, SD = 18.7; 4 left-handers: mean EHI score = -66.3, SD = 34.9; 2 ambidextrous: mean EHI score = 31.0, SD = 43.8) performed two successive reaching movements with either hand to one of two target locations. Each trial started with the presentation of the start locations in grey (RGB [128 128 128]) at the bottom of the screen. Participants then moved and maintained the cursors in these locations for 500 ms, causing the start locations to disappear. Then, the prime target appeared at the left or right target location in one of the four colors: orange (RGB [255 128 0], teal (RGB [0 255 128]), blue (RGB [0 128 255], or violet (RGB [128 0 255]), and participants moved the cursor of the respective arm to the target location. The color indicated which hand to use. To avoid perceptual repetition effects, different colors were used for the prime and probe movements, and color-hand assignment was randomized across participants. After the cursor was maintained at the location for 250 ms, the Kinarm moved the arm back to the start location. The back-transport took about 1500 ms and followed a bell-shaped velocity profile. After the cursor was maintained at the start location for 500ms, the probe target (left or right location) appeared (again in one of the four colors) and participants moved the cursor of the respective hand to the target location. After the cursor was maintained at the location for 250 ms, the Kinarm then moved the arm again back to the start location, followed by a 1000 ms intertrial interval. As in OA of Experiment Set 1, 50% of the trials were stop trials. During stop trials, the prime target location turned red (stop-signal, RGB [255 0 0]) 300 ms after prime target onset for 500 ms, signaling participants to withhold their movement (we used a larger delay for the stop signal in the HS task to acknowledge that this task required the additional process of effector selection^46^). Participants had to maintain the cursor within the start location for 2000 ms (to approximately match the duration between prime target onset and probe target onset of go trials), before the probe phase was initiated. During go trials, participants had to reach the target within 1200 ms after target onset, otherwise the message “Too slow!” was shown on the screen in red for 500 ms. If participants moved the wrong cursor out of the start position, the message “Wrong hand!” was displayed on the screen in red for 500 ms. If participants moved the cursor out of the start location during a stop trial, the message “Don’t move!” was displayed on the screen in red for 500 ms. Participants performed 13 blocks, each consisting of 64 trials. Half of the trials were go-trials, the other half stop-trials.

Movements were made to one of the two target locations with either the left or the right hand during the prime and probe, yielding 16 different prime-probe combinations (i.e., 2 prime hand x 2 prime target location x 2 probe hand x 2 probe target location), and hence a total of 32 different conditions. In each block, each condition was presented twice, and the order of presentation was randomized. The first block was considered practice and not analyzed. In total, the experiment comprised 768 trials (without practice trials) and took about 1.5 hours. We re-coded conditions depending on whether the hand was repeated for prime and probe (factor Hand: Repeat, Switch), and whether movement direction was repeated for prime and probe (factor Movement Direction: Repeat, Switch). We coded movement direction egocentrically, such that a prime movement with the left hand to the (contralateral) right target followed by probe movement with the right hand to the (contralateral) left target would be considered a movement direction repeat trial.

#### Experiment Set 2: Mechanical Stop

##### Obstacle Avoidance

30 participants (15 female, 15 male; mean age = 25.0 years, SD = 2.4; 25 right-handers: mean EHI score = 92.2, SD = 16.8, 4 left-handers: mean EHI score = -81.6, SD = 21.7, 1 ambidextrous: EHI = 47.4) controlled a blue circular cursor (0.5 cm diameter) to perform two center-out reaching movements in succession. The procedure was identical to the OA task in Experiment Set 1 with the following exceptions: Move trials were identical to go trials of Experiment Set 1. During block trials, the target (and obstacle, if present) during the prime phase was displayed in white, but movement execution was physically constrained by mechanical channels generated by the Kinarm. We implemented four force channels (channel width = 0.5 cm, wall width = 40000 cm, wall stiffness = 4000 N/m, damping of 10 Ns/m) each connecting the two target locations which were located diametral on the imaginary circle (e.g., one force channel between the 22.5° and the 202.5° target locations). These channels constrained movements during the prime phase of stop trials to a 0.5 x 0.5 cm square centered on the central start location. In these trials, the cursor had to maintain in the start location for 1000 ms before the probe phase commenced, and participants were instructed not to try to break out of force channels. Further, mechanical feedback was provided when the cursor hit the obstacle. During move trials, participants had to move the cursor into the target position within 1000 ms after target onset, otherwise the message “React faster!” was shown on the screen in red for 1000 ms. If participant hit the obstacle, it turned red and the message “You shall not pass!” was displayed on the screen in red for 1000 ms. Participants completed 640 trials split in 10 blocks preceded by one practice block that was not analyzed.

##### Hand Selection

35 individuals (13 female, 22 male; mean age = 26.3 years, SD = 8.5; 31 right-handers: mean EHI score = 95.0, SD = 7.7; 3 left-handers: mean EHI-score = -57.7, SD = 62.7; 1 ambidextrous: EHI-score = 20.0) participated. The procedure was identical to Experiment Set 1 with the following exceptions: During stop trials, following prime target presentation, movement execution of both arms was physically constrained (as in Experiment Set 2 OA task). We implemented three force channels (channel width = 0.5 cm, wall width = 40000 cm, wall stiffness = 4000 N/m, damping of 10 Ns/m), one connecting the two start locations, and one connecting the start location of each hand with the target, respectively. These channels constrained movements of each hand during the prime phase of stop trials to a 0.5 x 0.5 cm square centered on the respective start location. In these trials, the cursors had to maintain in the start locations for 2000 ms before the probe phase commenced, and participants were instructed not to try to break out of force channels. In total, participants completed 768 trials split in 12 blocks plus one practice block that was not analyzed.

#### Experiment Set 3: Passive Movements and Vision Control

##### Obstacle Avoidance

30 participants (23 female, 6 male, 1 diverse; mean age = 22.4 years, SD = 4.4; 28 right-handers: mean EHI score = 87.4, SD = 13.3, 2 left-handers: mean EHI score = -95.0, SD = 7.1) controlled a white circular cursor (0.5 cm diameter) to perform two center-out reaching movements in succession. The procedure was identical to Experiment Set 2 with the following exceptions: On a subset of trials (passive trials), the Kinarm robot moved the handle during the prime phase (from start to target and back to start) while participants holding onto it by replaying the participants’ movements recorded from active prime trials. We used two types of passive trials: during passive vision trials, the prime movement was performed by the Kinarm while target location and obstacle (if present) were visually displayed. The targets were displayed in blue (RGB [0 128 255]) or orange (RGB [255 128 0]), indicating if the movement had to be actively performed or was passively performed by the Kinarm (color-trial type assignment randomized across participants). During passive no visual feedback trials, the prime movement was performed also by the Kinarm but cursor, target, and obstacle were not visible.

Participants performed 10 experimental blocks, each consisting of 96 trials. Each block commenced with a sub-block of 32 active trials (each combination of 2 Obstacle Prime x 2 Obstacle Probe x 8 Target Location was performed once) during which we recorded participants prime movements followed by two sub-blocks of 32 passive trials during which the recorded prime movements were passively executed by the Kinarm. The first session (block 1-5) contained passive no feedback trials, the second session (block 6-10) contained passive feedback trials. Participants performed one practice block with 32 active and 32 passive trials prior to each feedback session. Thus, participants performed a total of 1088 trials (10 blocks with each 96 trials and 2 training blocks with 64 trials each).

##### Obstacle Avoidance Active movements with and without visual feedback

30 individuals (27 female, 3 male; mean age = 21.5 years, SD = 1.8; 24 right-handers: mean EHI score = 93.7, SD = 8.7; 6 left-handers: mean EHI-score = -85.8, SD = 5.6) participated. Task and procedure was identical to the OA task in Experiment Set 2 with the following exceptions: 250 ms after prime target (and obstacle, if present) appearance, the start location turned green signaling participants to initiate their movement (i.e., go cue; movements initiated prior the go-cue were considered errors and the respective message “Too early!” was displayed). In half of the trials, cursor, target, and obstacle disappeared concurrently with onset of the go cue (no vision trials). In the other half, they remained visible throughout the prime phase (vision trials). To ensure that participants would hit the target in no vision trials, we increased the logical size of the targets to 3 cm diameter (visual diameter remained identical, 1 cm). Participants completed 640 trials split in 10 blocks preceded by one practice block that was not analyzed.

##### Hand Selection Passive Movement Task

28 right-handed individuals (15 female, 13 male; mean age = 23.8 years, SD = 3.0; mean EHI-score = 98.4, SD = 4.7) participated. The procedure was identical to the HS task in Experiment Set 2 with the following exceptions: On a subset of trials (passive trials), the Kinarm robot moved the handle from the start to the target location during the prime phase while participants holding onto it by replaying the participants’ movements recorded from active prime trials. In some blocks, targets were visible during the passive movements (passive vision) and displayed in the same color as during the active prime trials. In some blocks, targets were not visually displayed during passive movement (passive no vision). Further minor changes were that the back-transport after prime and probe phase took about 2000 ms and followed a bell-shaped velocity profile, and participants had to reach the target within 2000 ms after target onset. Participants performed 4 blocks, each consisting of 192 trials. Each block commenced with a sub-block of 64 active trials followed by two sub-blocks each with 64 passive trials either with or without visual feedback.

Passive visual feedback and passive no visual feedback blocks were performed in alternation with the order counterbalanced across participants. In each sub-block, all 16 prime-probe combinations (i.e., 2 prime hand x 2 prime target location x 2 probe hand x 2 probe target location) were repeated four times in a randomized order. Thus, participants performed 256 active and 256 passive visual feedback, and 256 passive no feedback trials, yielding a total of 768 trials, preceded by a short practice block to familiarize themselves with the color-hand assignment and the passive arm displacement. The experiment took about three hours.

#### Experiment Set 4: No-Go

##### Obstacle Avoidance

We analyzed data from 30 right-handed participants (18 female, 12 male; mean age = 24.4 years, SD = 6.1; mean EHI score = 90.0, SD = 14.0). The experimental procedures were identical to the OA task in Experiment Set 1, except that during no-go trials, the prime target location was immediately shown in red (instead of the 150ms delay), signaling participants to withhold their movement. Participants completed 640 trials split in 10 blocks preceded by one practice block that was not analyzed.

##### Obstacle Avoidance with trial types in separate blocks

30 participants (21 female, 9 male; mean age = 22.4, SD = 3.2; 28 right-handers: mean EHI score = 72.2, SD = 21.1, 2 left-handers: mean EHI score = -70.7, SD = 11.6) completed this task. The experiment was identical to the previous experiment except that no-go trials and go trials were presented in separate blocks. Half of the participants performed 10 blocks (plus one practice block) with only no-go trials (32 trials per block) followed by 10 blocks (plus one practice block) with only go trials (32 trials per block), yielding a total of 640 trials plus 64 practice trials. For the other half of the participants, the order was reversed.

##### Hand Selection

29 individuals (24 female, 1 male; mean age = 24.1years, SD = 5.5; 25 right-handers: mean EHI score = 86.3, SD = 15.5, 3 left-handers: mean EHI score = -90.0, SD = 17.3, 1 ambidextrous: EHI score = 0) articipated. The procedure was identical to the HS task in Experiment Set 1, except that a red circle (1.2 cm in diameter, stop signal) was displayed in the center of and concurrently with the prime target cue, signaling participants to withhold their movement.

### Data processing and analysis

We processed kinematic data with custom-written scripts in MATLAB (version R2023a; The MathWorks, Natick, MA). We filtered the data from the mouse tracking experiments using a second-order butterworth filter with a cutoff frequency of 10 Hz and the Kinarm data using a third-order zero-lag double-pass filter with a cutoff frequency of 10 Hz. We determined reach onset as the time of the sample in which the vectorial velocity exceeded 50 mm/ s and reach offset as the time of the sample in which the cursor entered the target location of the respective phase (i.e., prime and probe). For the OA task, we used initial reach error as a measure of feedforward effects and we quantified history effects as the difference in initial reach error between probe movements for which the prime movement had, vs. had not, involved an obstacle (i.e., initial reach bias). For the mousing tracking experiments, we defined initial reach error as the absolute value of the angular deviation between the vector connecting cursor position at reach onset and the position of the cursor when it left the start location and the vector connecting cursor position at reach onset and the target center. The average time between movement onset and the timepoint when the cursor left the start location was considerably below the time necessary to utilize visual feedback (∼100ms, Scott, 2016; OA task Experiment Set 1: prime movement mean = 36 ms, SD = 10, probe movement mean = 43 ms, SD = 11; OA task Experiment Set 4: prime movement mean = 36 ms, SD = 10, probe movement mean = 42 ms, SD = 12). For the Kinarm obstacle avoidance task experiments, we defined initial reach error as the absolute value of the angular deviation between the vector connecting cursor position at reach onset and the position of the cursor 50 ms later and the vector connecting cursor position at reach onset and the target center. For the HS task, we used reaction time (RT) alongside initial reach error as a measure of feedforward effects. We defined reaction time (RT) as the time between target onset and reach onset of probe movements. We quantified history effects as the difference in RT between trials for which 1) different vs the same hand was used for prime and probe movements (hand repetition effect, HRE) and 2) movements were performed in different vs same directions (coded in an egocentric reference frame; movement repetition effect, MRE). For the HS task, we defined initial reach error as the signed value of the angular deviation between the vector connecting cursor position at reach onset and the position of the cursor 50 ms later and the vector connecting cursor position at reach onset and the target center. We mirrored the values for the left hand such that positive values indicate an inward bias. We quantified history effects (i.e., initial reach bias) as difference in initial reach error between probe movements for diagonal vs. straight prime movements, positive values indicate that probe movements were more inclined towards the body-midline following diagonal vs. straight prime movements.

We used three measures to consider feedback-related effects: First, as a measure of reach accuracy, we obtained reach endpoints as the cursor position 150ms after it entered the target and when its resultant velocity was below a threshold (5cm/s and 10cm/s for the OA and HS task, respectively) and calculated absolute reach endpoint error as the Euclidean distance (in mm) of the reach endpoints to the target center. Second, as a measure of reach precision, we calculated the area of the 95% confidence ellipse areas (in mm^2^) for these reach endpoints. We quantified history effects as the difference in absolute reach endpoint errors/ ellipse areas between trials with different versus same prime– probe movement requirements (OA: obstacle presence/ absence; HS: different vs. same movement direction).

Positive values indicate better performance (i.e., smaller endpoint errors/ ellipse areas) for same prime–probe movement requirements. We chose the combined duration/velocity criterion to minimize any potential conflating influence of feedforward processes and/ or speed-accuracy tradeoffs on terminal feedback. Third, we quantified final reach error as the absolute value of the angular deviation between the vector connecting cursor position at reach onset and the target center and the vector connecting cursor position at reach onset and target hit.

### Data exclusion

Passive trials in which an error occurred during the corresponding active trials were skipped (Experiment Set 3 OA task: N = 1328; HS task: N = 1286). Further, we excluded trials in which participants moved the cursor out of the start location during a stop-signal/no-go trial (Experiment Set 1 OA task: N = 2302, 24% of stop-signal trials; HS task: N = 3499, 23.4% of stop trials; Experiment Set 4 OA task: N = 358, 3.7% of no-go trials; OA control task: N = 349, 3.6% of no-go trials; HS task: N = 85, 0.8% of no-go trials), trials in which they reacted to slowly during the prime (Experiment Set 1 OA task: N =153; HS task: N = 191; Experiment Set 2 OA task: N = 815; HS task: N = 59; Experiment Set 3 OA task: N = 1107; OA control task: N = 696; HS task: N = 46; Experiment Set 4 OA task: N = 47; OA control task: N = 18; HS task: N = 88) or probe phase (Experiment Set 1 OA task: N =174; HS task: N = 177; Experiment Set 2 OA task: N = 987; HS task: N = 55; Experiment Set 3 OA task: N = 900; OA control task: N = 69; HS task: N = 72; Experiment Set 4 OA task: N = 81; OA control task: N = 134; HS task: N = 131), trials in which they hit the obstacle or used the wrong hand during the prime (Experiment Set 1 OA task: N =127; HS task: N = 861; Experiment Set 2 OA task: N = 113; HS task: N = 1458; Experiment Set 3 OA task: N = 77; OA control task: N = 154; HS task: N = 290; Experiment Set 4 OA task: N = 89; OA control task: N = 183; HS task: N = 970) or probe phase (Experiment Set 1 OA task: N =230; HS task: N = 1365; Experiment Set 2 OA task: N = 339; HS task: N = 1405; Experiment Set 3 OA task: N = 118; OA control task: N = 156; HS task: N = 864; Experiment Set 4 OA task: N = 264; OA control task: N = 294; HS task: N = 1316), trials in which movement onset could not be detected during the prime (Experiment Set 1 OA task: N =149; HS task: N = 0; Experiment Set 2 OA task: N = 0; HS task: N = 0; Experiment Set 3 OA task: N = 0; OA control task: N = 0; HS task: N = 0; Experiment Set 4 OA task: N = 144; OA control task: N = 272; HS task: N = 0) or probe phase (Experiment Set 1 OA task: N =166; HS task: N = 27; Experiment Set 2 OA task: N = 0; HS task: N = 15; Experiment Set 3 OA task: N = 0; OA control task: N = 0; HS task: N = 0; Experiment Set 4 OA task: N = 178; OA control task: N = 258; HS task: N = 19), and trials which did not exhibit smooth trajectories (i.e., movements with more than two velocity peaks) and trials in which participants exerted force in movement direction during passive trials (Experiment Set 1 OA task: N =768; HS task: N = 0; Experiment Set 2 OA task: N = 238; HS task: N = 0; Experiment Set 3 OA task: N = 667; OA control task: N = 10; HS task: N = 175; Experiment Set 4 OA task: N = 666; OA control task: N = 683; HS task: N = 0). This led to exclusion of 3851 trials (20.1%) in Experiment Set 1 OA task, 5761 trials (19.3%) in Experiment Set 1 HS task, 2415 trials (12.7%) in Experiment Set 2 OA task, 2843 trials (10.6%) in Experiment Set 2 HS task, 4196 trials (14.6%) in Experiment Set 3 OA task, 1338 (7.0%) in Experiment Set 3 OA control task, 2733 trials (12.7%) in Experiment Set 3 HS task, 1756 (9.1%) in Experiment Set 4 OA task, 2084 trials (10.9%) in Experiment Set 4 OA control task, and 2543 trials (11.0%) in Experiment Set 4 HS task. Finally, we removed trials in which RT was shorter than 100 ms (obstacle avoidance task)/ 150 ms (hand selection task) or longer than 1200 ms and trials in which MT was longer than 1000 ms (Experiment Set 1 OA task: N =187; HS task: N = 0; Experiment Set 2 OA task: N = 210; HS task: N = 0; Experiment Set 3 OA task: N = 699; OA control task: N = 19; HS task: N = 146; Experiment Set 4 OA task: N = 268; OA control task: N = 626; HS task: N = 5).

### General Statistical approach

To analyze our data, we fitted Bayesian regression models created in Stan (http://mc-stan.org/) and accessed with the package brms version 2.23^47^ in R 4.5.1^48^. We fit separate models for each of our dependent variables: Initial reach error of probe movement, RT, final reach error, end point precision, and end point accuracy. We set orthogonal contrasts using the set_sum_contrast() command in afex version 1.5-1^49^ and included random intercepts and slopes for all main effects and interactions. We used normal, skewed-normal, and (shifted) log-normal distributions to estimate parameters and specified uninformative priors for population-level (i.e., fixed) effects (see specific models). The estimation of parameters’ posterior distributions was obtained by Hamiltonian Monte-Carlo sampling with 4 chains, 1,000 sample warmup, and 11,000 iterations and checked visually for convergence (high ESS and Rhat ≈ 1). We used the package bayestestR version 0.18.1^50,51^ to describe the parameters of our models. We report the median as a point estimate of centrality and the 95% credible interval (CI) computed based on the highest-density interval (HDI) to characterize the uncertainty related to the estimation. Following current guidelines for Bayesian inference^50–52^, we distinguish between the *existence* and the *significance* of an effect, as these index conceptually different questions and need not agree. Existence — whether an effect is credibly different from zero in a given direction — was assessed using the Probability of Direction (pd), the proportion of the posterior distribution that shares the sign of the median.

Following the reference points proposed by Makowski et al.^50^, we interpret pd ≤ 95% as *uncertain*, pd > 95% as *possibly existing*, pd > 97% as *likely existing*, and pd > 99% as *probably existing*. Significance — whether an effect is large enough to be considered non-negligible for practical purposes — was assessed by calculating the percentage of the HDI falling inside a region of practical equivalence (ROPE) of ±0.1 effect sizes around zero^52^. If the HDI fell completely outside the ROPE, the effect was considered significant (i.e., decidedly non-negligible); if the ROPE fully contained the HDI, the effect was considered negligible (i.e., practically equivalent to zero); partial overlap was treated as inconclusive with respect to significance. Because pd and ROPE address different questions, an effect can exist (high pd) while remaining too small to be considered significant (substantial or complete ROPE overlap); we report both indices for every effect and interpret them jointly rather than treating either in isolation as a binary “present/absent” verdict.

### Specific models

#### Obstacle avoidance task

For initial reach error and final reach error, we fit separate Bayesian regression models depending on whether an obstacle was present or absent during the probe phase, since the distributions were vastly different between these conditions. For both models, we included the within-subject categorical variables prime phase (obstacle, no obstacle) and trial type (Experiment Set 1: go, stop; Experiment Set 2: move, block; Experiment Set 3: active, passive vision, passive no vision; Experiment Set 4: go, no-go). We modeled Initial reach error using truncated normal and log-normal distributions (lower bound [lb]: 0°; upper bound [ub]180°) for trials with and without an obstacle in the probe phase, respectively. We modeled Final reach error using skewed normal and truncated (lb: 0°, ub: 4°) normal distributions for trials with and without an obstacle in the probe phase, respectively. For end point variability (i.e., precision) and end point error (i.e., accuracy), we also included the within-subject categorical variables prime phase (obstacle, no obstacle), and we used truncated (precision: lb: 0mm^2^; accuracy: lb: 0mm, ub: 5mm) normal distributions.

#### Hand selection task

For RT, we included the within-subject categorical variables hand repetition (repeat, switch), movement repetition (repeat, switch), trial type (Experiment Set 1: go, stop; Experiment Set 2: move, block; Experiment Set 3: active, passive vision, passive no vision; Experiment Set 4: go, no-go), and all interactions as independent predictors. We modeled RT using shifted log-normal distributions. For initial reach error and final reach error, we fit separate models with the within-subject categorical variables hand (repeat, switch), trial type (Experiment Set 1: go, stop; Experiment Set 2: move, block; Experiment Set 3: active, passive vision, passive no vision; Experiment Set 4: go, no-go), and prime movement direction (straight, diagonal), and probe movement direction (straight, diagonal), using normal distributions (initial reach error) and truncated log-normal (lb: 0°, ub: 10°) distributions (final reach error). For end point variability (i.e., precision) and end point error (i.e., accuracy), we included the within-subject categorical variables hand repetition (repeat, switch), movement repetition (repeat, switch), trial type (Experiment Set 1: go, stop; Experiment Set 2: move, block; Experiment Set 3: active, passive vision, passive no vision; Experiment Set 4: go, no-go), and we used log-normal (precision) and truncated log-normal distributions (accuracy: ub: 20 mm). Exact model specifications for each model can be obtained from the code available in the corresponding authors’ osf repository (https://osf.io/xhvm7/overview).

## Data availability

Data and code supporting the findings of the present article will me made publicly available via the Open Science Framework website (https://osf.io/xhvm7/overview).

## Acknowledgments

We thank Evelyn Augustin, Tobias Dietrich, Elena Franchi, Linda Gottfried, Hannah Haslberger, Juliana Möller, Arne Morfeld, Roman Topmöller, Étienne Wilker, and Niklas Wolberg for their help with data collection.

## Author contributions

C. S. and T.H. conceived and designed research; C. S. performed experiments; C. S. analyzed data; C. S. and T.H. interpreted results of experiments; C. S. prepared figures; C. S. drafted manuscript; C. S. and T.H. edited and revised manuscript; C. S. and T.H. approved final version of manuscript.

