## Supplementary Materials for "Motor planning and execution establish distinct feedforward and feedback motor histories"

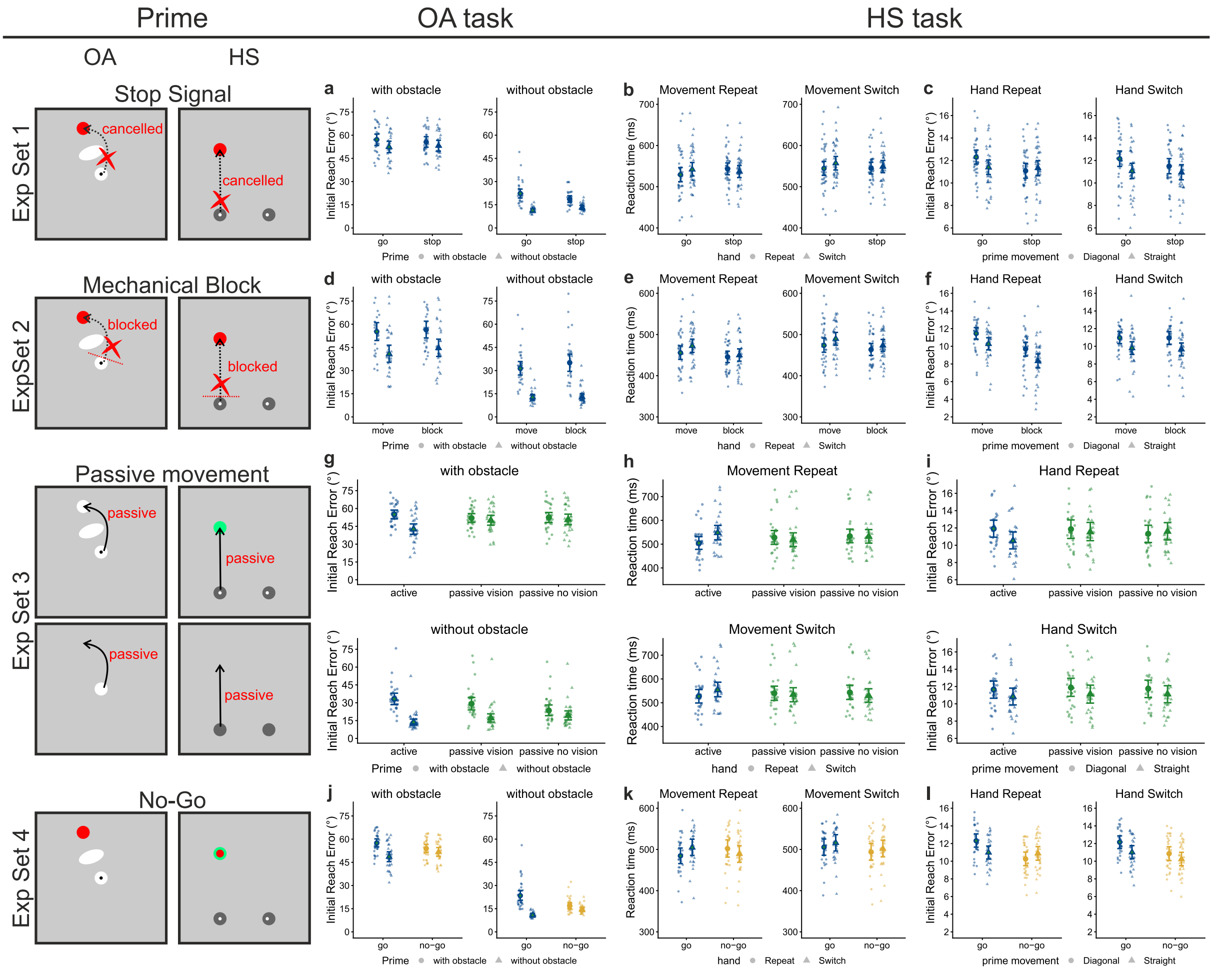


**Supplementary Figure 1.** Motor history effects in feedforward measures. Estimated marginal means for all experiments. For the obstacle avoidance (OA) task, we calculated initial reach error as a function of prime phase (with obstacle, without obstacle), probe phase (with obstacle, without obstacle), and trial type (Exp. 1: go, stop; Exp 2.: move, block; Exp. 3a: active, passive vision, passive no vision; Exp. 3b: vision, no vision; Exp.4ab: go, no-go). For the hand selection (HS) task, we calculated reaction time (RT) as a function of hand (repeat, switch), movement (repeat, switch), and trial type (as in OA), and initial reach error as a function of hand (repeat, switch), prime movement (straight, diagonal), and trial type (as in OA). Large symbols represent the group estimates and error bars the 95% highest-density interval (HDI). Small symbols represent individual participant estimates.

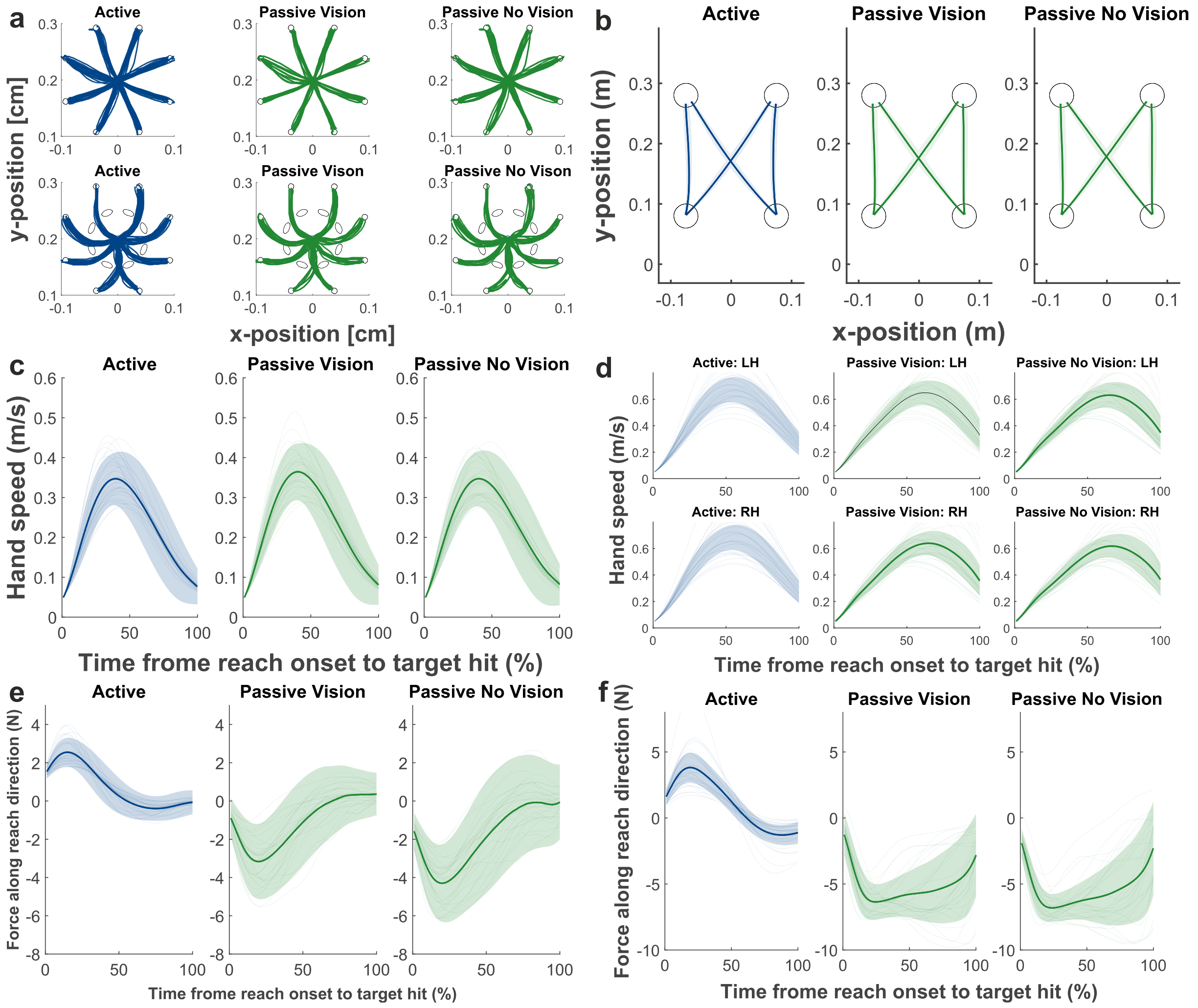


**Supplementary Figure 2.** In Experiment Set 3, the prime consisted of robot-imposed, passive movements that replayed each participant's previously recorded, active trajectories. Active and passive prime movements exhibited highly similar spatiotemporal characteristics (**a-d**), and participants did not, against the instructions to be guided passively, exert force in movement direction (**ef**). Movement trajectories from an exemplary participant in the OA task (**a**), movement trajectories in the HS task (**b**), hand speed profiles (**cd**), and applied forces (**ef**) for the OA and HS task, respectively. Solid lines represent mean values, and shaded areas represent ±1 standard deviations.

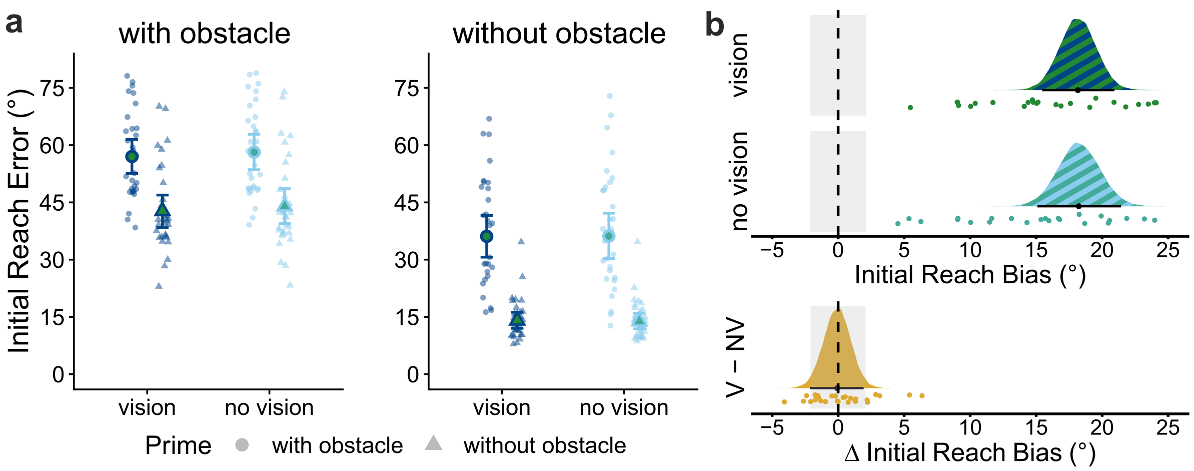


**Supplementary Figure 3.** Motor history effects in feedforward measures are similar for active prime movements performed with or without visual display of cursor and target configuration. **A** Initial reach error as a function of prime phase (with obstacle, without obstacle), probe phase (with obstacle, without obstacle), and trial type (vision, no vision). Large symbols represent the group estimates and error bars the 95% highest-density interval (HDI). Small symbols represent individual participant estimates. **B** Initial reach bias, calculated as difference of initial reach error in trials with obstacle minus without an obstacle in the prime phase for vision (V) and no vision (NV) trials, and difference in initial reach bias between vision and no vision trials (initial reach bias vision: 18.2° [15.4, 20.9], pd = 100%, 0% in ROPE, no vision: 18.2° [15.1, 21.4], pd = 100%, 0% in ROPE, difference vision vs no vision: -0.1° [-2.1, 1.9], pd = 52.68%, 99% in ROPE). Colored areas: posterior distributions. Black dots: median; error bars: 95% highest-density interval (HDI); grey shaded areas: region of practical equivalence (ROPE). V = vision, NV = no vision.

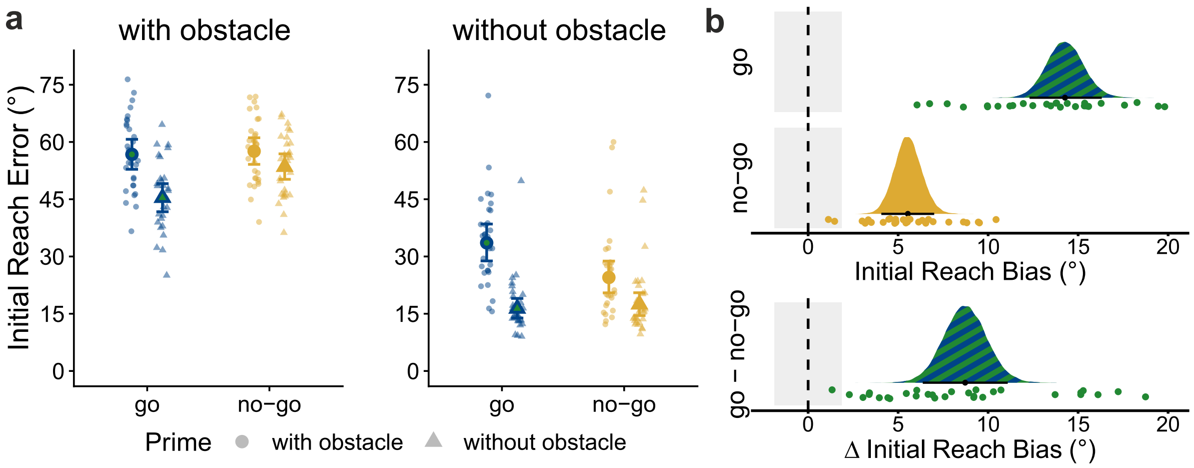


**Supplementary Figure 4.** Residual motor history effects in feedforward measures persist in no-go trials when presented in separate blocks. **A** Initial reach error as a function of prime phase (with obstacle, without obstacle), probe phase (with obstacle, without obstacle), and trial type (go, no-go). Large symbols represent the group estimates and error bars the 95% highest-density interval (HDI). Small symbols represent individual participant estimates. **B** Initial reach bias, calculated as difference of initial reach error in trials with obstacle minus without an obstacle in the prime phase for go and no-go trials, and difference in initial reach bias between go and no-go trials (initial reach bias go: 14.3° [12.3, 16.3], pd = 100%, 0% in ROPE, no-go: 5.5° [4.1, 7.0], pd = 100%, 0% in ROPE, difference go vs no-go: 9.7° [6.4, 11.1], pd = 100%, 0% in ROPE). Colored areas: posterior distributions. Black dots: median; error bars: 95% highest-density interval (HDI); grey shaded areas: region of practical equivalence (ROPE).

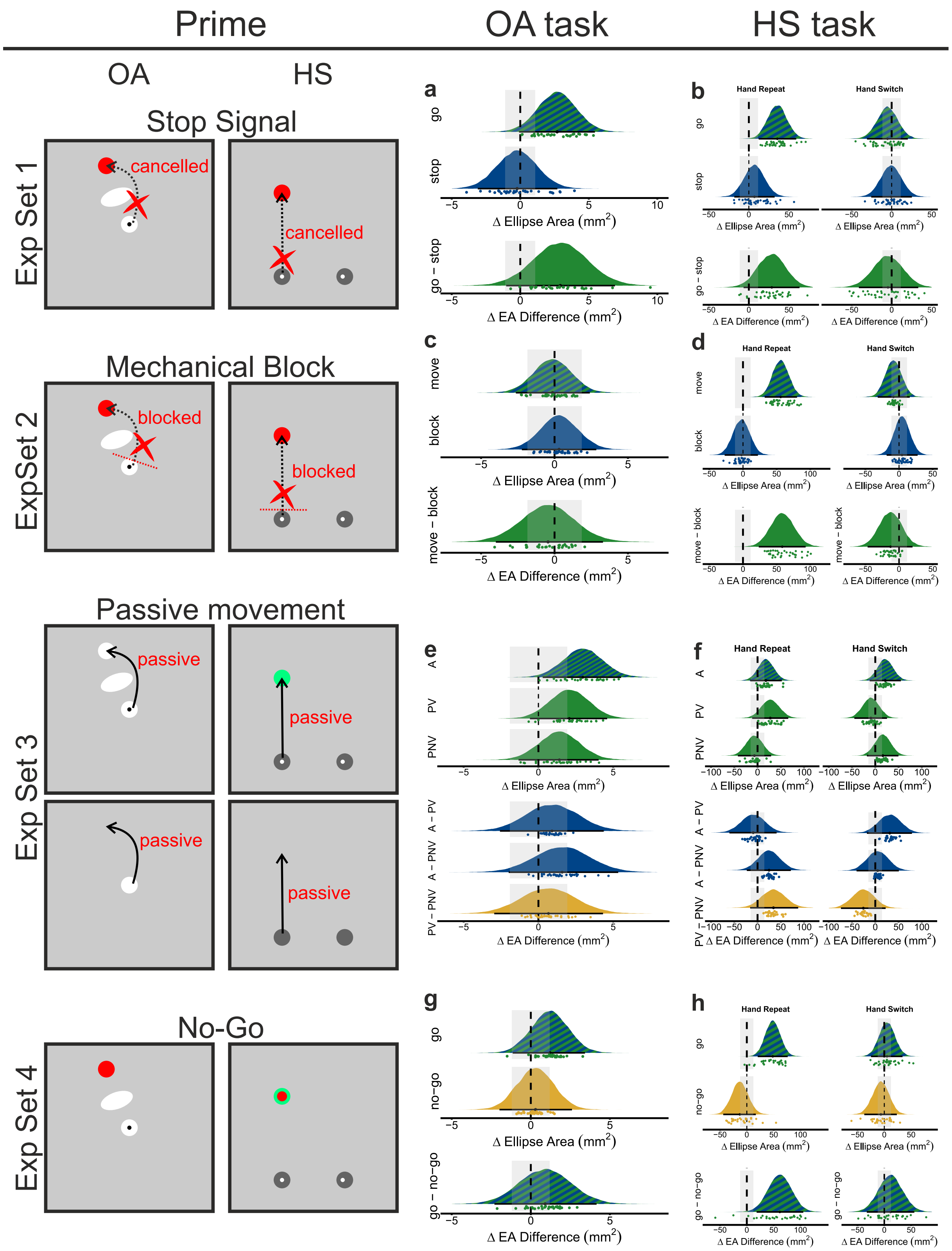


**Supplementary Figure 5.** Motor history effects in reach endpoint precision, quantified as the difference of the 95% confidence ellipse areas of reach endpoints between trials with different versus same prime–probe movement requirements (OA: obstacle presence/ absence; HS: different vs. same movement direction). Positive values indicate better performance. Colored areas: posterior distributions. Black dots: median; error bars: 95% highest-density interval (HDI); grey shaded areas: region of practical equivalence (ROPE). In the OA task, prime movements that were planned and executed produced a small (but negligible in magnitude) difference in probe reach precision (pooled across the four experiments: 1.59mm² [-0.21, 3.35], pd = 96%, 60% in ROPE. When execution was cancelled via a delayed stop-signal, there was no difference (Exp. 1: -0.23mm² [-3.10, 2.70], pd =

56%, 37% in ROPE), nor when execution was mechanically blocked while the plan remained intact (Exp. 2: 0.31mm² [-2.22, 2.85], pd = 60%, 73% in ROPE). Likewise, there was no difference following robot-imposed passive primes, regardless of whether the visual display was shown (vision: 2.07mm² [-0.57, 4.59], pd = 94%, 48% in ROPE; no vision: 1.40mm² [-1.33, 4.03], pd = 85%, 61% in ROPE), nor following primes with neither planning nor execution (Exp. 4: 0.30mm² [-1.98, 2.60], pd = 71%, 37% in ROPE). In the HS task, there was an improvement in reach precision for matched prime–probe movement requirements during active trials when the same hand was used across prime and probe (pooled across the four experiments: 40.24mm² [24.50, 55.50], pd > 99%, 0% in ROPE), but not when hands were switched (2.45mm² [-13.4, 18.4], pd = 62%, 76% in ROPE). Cancelling a planned movement after a stop-signal (Exp. 1) or mechanical blocking (Exp. 2) revealed no difference (Exp. 1: hand repeat: 6.98mm² [-18.14, 32.26], pd = 71%, 45% in ROPE; hand switch: -0.68mm² [-25.56, 25.26], pd = 52%, 45% in ROPE; Exp. 2: hand repeat: -2.33mm² [-27.20, 22.20], pd = 58%, 46% in ROPE; hand switch: 4.39mm² [-18.40, 27.60], pd = 65%, 50% in ROPE). Likewise, there was no difference for passive primes (Exp. 3), regardless of whether vision was present (hand repeat: 27.31mm² [-11.56, 65.96], pd = 92%, 34% in ROPE; hand switch: -9.50mm² [-44.93, 25.22], pd = 71%, 41% in ROPE), or not (hand repeat: -6.81mm² [-42.28, 29.10], pd = 65%, 41% in ROPE; hand switch: 16.28mm² [-18.00, 49.85], pd = 83%, 43% in ROPE), nor when both planning and execution were absent (Exp. 4: hand repeat: -13.06mm² [-43.90, 16.80], pd = 80%, 41% in ROPE; hand switch: -6.37mm² [-37.30, 23.30], pd = 66%, 41% in ROPE).

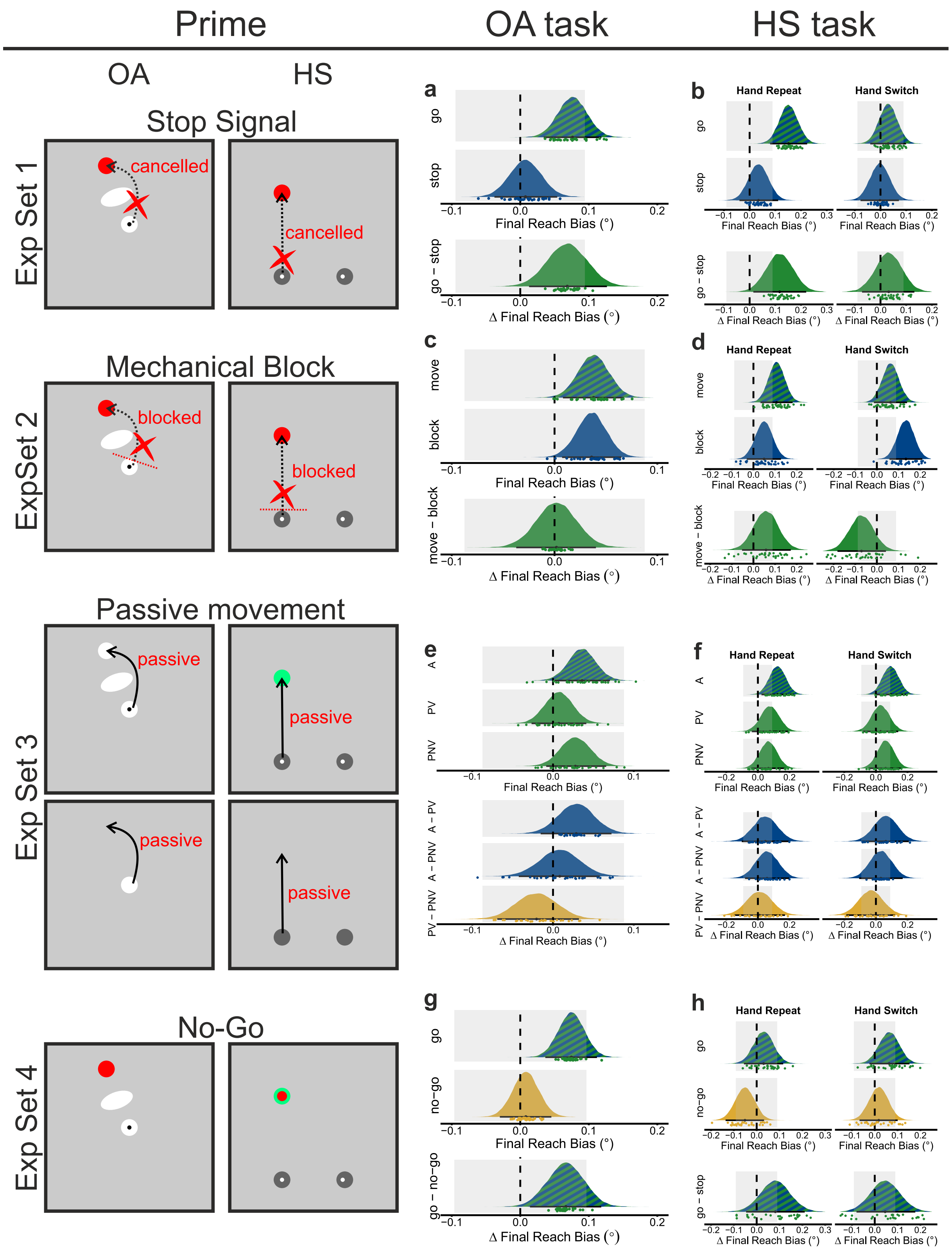


**Supplementary Figure 6.** Motor history effects in final reach error (i.e., final reach bias), quantified as the difference in final reach error between trials with different versus same prime–probe movement requirements (OA: obstacle presence/ absence; HS: different vs. same movement direction). Positive values indicate better performance. Colored areas: posterior distributions. Black dots: median; error bars: 95% highest-density interval (HDI); grey shaded areas: region of practical equivalence (ROPE). In the OA task, prime movements that were planned and executed produced a small but reliable final reach bias in the probe (Exp. 1: 0.08° [0.03, 0.12], pd > 99%, 72% in ROPE; Exp. 2: 0.04° [0.01, 0.07], pd > 99%, 100% in ROPE; Exp. 3: 0.04° [0.00, 0.07], pd = 98%, 100% in ROPE; Exp. 4: 0.07° [0.04, 0.11], pd Y 99%, 79% in ROPE). When planning was present, but execution was cancelled via a delayed stop-signal (Exp. 1), the final reach bias was absent (0.01° [-0.04, 0.05], pd = 63%, 100% in ROPE). When planning was present and execution was mechanically blocked, a final reach bias was evident (Exp. 2: 0.04° [0.01, 0.07], pd > 99%, 100% in ROPE). Robot-imposed passive primes (Exp. 3) did not evoke a final reach bias, regardless of whether the visual display was shown or not (vision: 0.01° [-0.03, 0.04], pd = 66%, 100% in ROPE; no vision : 0.02° [-0.01, 0.07], pd = 93%, 100% in ROPE). Similarly, no final reach bias was observed when primes proceeded without both planning and execution (Exp. 4: 0.01° [-0.03, 0.05], pd = 67%, 100% in ROPE). In the HS task, the final reach bias during active trials was small but reliable when the same hand was used across prime and probe in three out of the four experiments (Exp. 1: 0.15° [0.08, 0.22], pd > 99%, 7% in ROPE; Exp. 2: 0.10° [0.03, 0.18], pd > 99%, 37% in ROPE; Exp. 3: 0.12° [0.02, 0.23], pd > 99%, 34% in ROPE; Exp. 4: 0.03° [-0.06, 0.12], pd = 76%, 85% in ROPE), but was absent when hands switched (Exp. 1: 0.03° [-0.04, 0.10], pd = 80%, 93% in ROPE; Exp. 2: 0.06° [-0.01, 0.14], pd = 95%, 66% in ROPE; Exp. 3: 0.09° [-0.01, 0.20], pd = 96%, 49% in ROPE; Exp. 4: 0.06° [-0.02, 0.15], pd = 92%, 65% in ROPE). As in OA, cancelling a planned movement after a stop-signal (Exp. 1) eliminated the final reach bias entirely (hand repeat: 0.03° [-0.04, 0.11], pd = 82%, 86% in ROPE; hand switch: 0.00° [-0.08, 0.07], pd = 52%, 100% in ROPE). Mechanical blocking (Exp. 2), however, produced a bias only on hand-switch trials (0.13° [0.06, 0.21], pd > 99%, 20% in ROPE), not on hand-repeat trials (0.05° [-0.03, 0.13], pd = 89%, 75% in ROPE). Passive primes did not produce a final bias regardless of visual display (with vision — hand repeat: 0.08° [-0.04, 0.19], pd = 91%, 56% in ROPE; hand switch: 0.03° [-0.08, 0.14], pd = 70%, 77% in ROPE; without vision — hand repeat: 0.06° [-0.04, 0.17], pd = 89%, 61% in ROPE; hand switch: 0.06° [-0.05, 0.16], pd = 88%, 65% in ROPE), and also not when both planning and execution were absent (Exp. 4: hand repeat: -0.05° [-0.14, 0.03], pd = 88%, 73% in ROPE; hand switch: 0.01° [-0.07, 0.10], pd = 66%, 94% in ROPE).

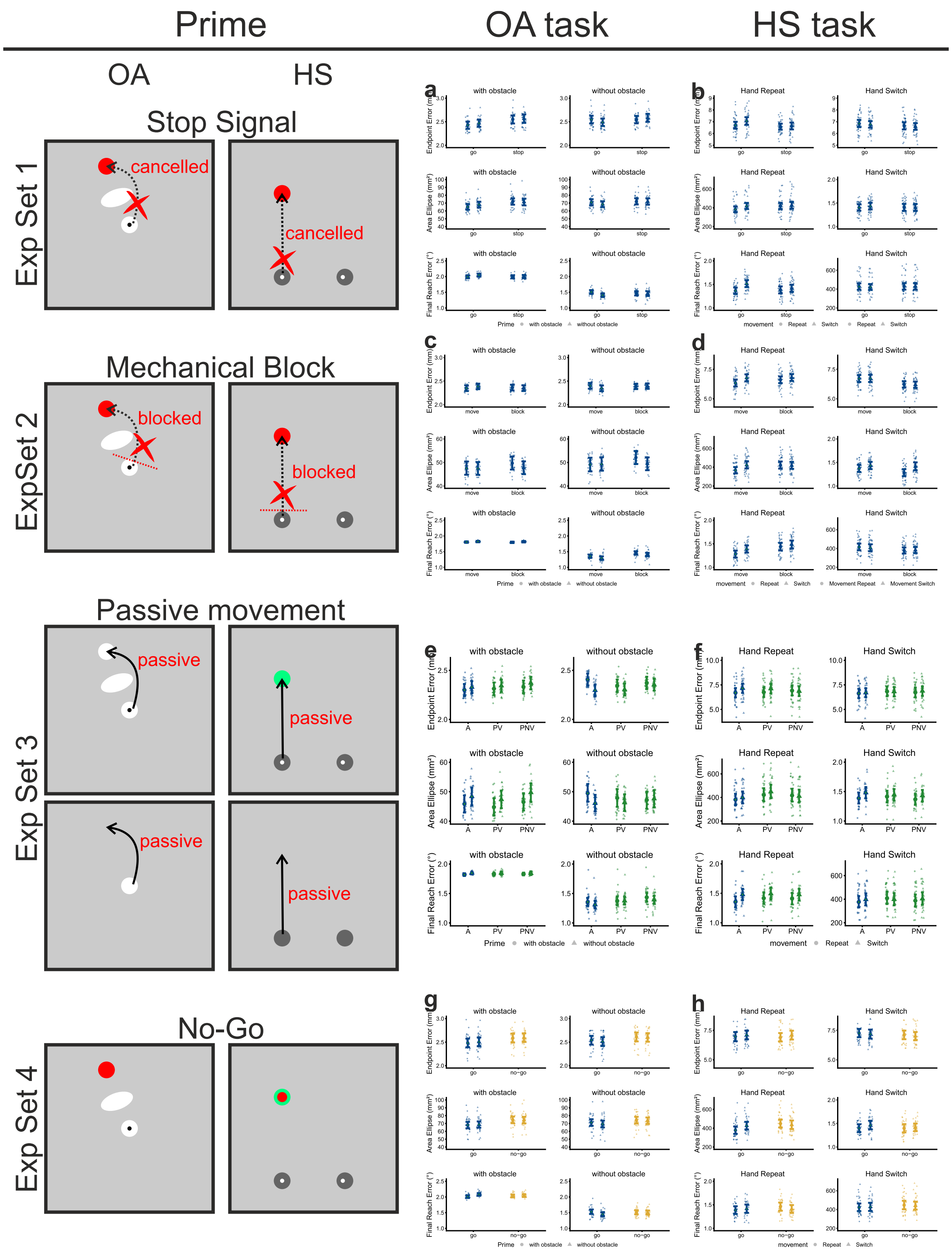


**Supplementary Figure 7.** Motor history effects in feedback measures. Estimated marginal means for all experiments. For the obstacle avoidance (OA) task, we calculated endpoint error, variability of endpoint error, and final reach error as a function of prime phase (with obstacle, without obstacle), probe phase (with obstacle, without obstacle), and trial type (Exp. 1: go, stop; Exp 2.: move, block; Exp. 3a: active, passive vision, passive no vision; Exp. 3b: vision, no vision; Exp.4ab: go, no-go). For the hand selection (HS) task, we calculated endpoint error, variability of endpoint error, and final reach error as a function of hand (repeat, switch), movement (repeat, switch), and trial type (as in OA), and initial reach error as a function of hand (repeat, switch), prime movement (straight, diagonal), and trial type (as in OA). Large symbols represent the group estimates and error bars the 95% highest-density interval (HDI). Small symbols represent individual participant estimates.
